# DeCOIL: Optimization of Degenerate Codon Libraries for Machine Learning-Assisted Protein Engineering

**DOI:** 10.1101/2023.05.11.540424

**Authors:** Jason Yang, Julie Ducharme, Kadina E. Johnston, Francesca-Zhoufan Li, Yisong Yue, Frances H. Arnold

## Abstract

With advances in machine learning (ML)-assisted protein engineering, models based on data, biophysics, and natural evolution are being used to propose informed libraries of protein variants to explore. Synthesizing these libraries for experimental screens is a major bottleneck, as the cost of obtaining large numbers of exact gene sequences is often prohibitive. Degenerate codon (DC) libraries are a cost-effective alternative for generating combinatorial mutagenesis libraries where mutations are targeted to a handful of amino acid sites. However, existing computational methods to optimize DC libraries to include desired protein variants are not well suited to design libraries for ML-assisted protein engineering. To address these drawbacks, we present DEgenerate Codon Optimization for Informed Libraries (DeCOIL), a generalized method which directly optimizes DC libraries to be useful for protein engineering: to sample protein variants that are likely to have both high fitness and high diversity in the sequence search space. Using computational simulations and wet-lab experiments, we demonstrate that DeCOIL is effective across two specific case studies, with potential to be applied to many other use cases. DeCOIL offers several advantages over existing methods, as it is direct, easy-to-use, generalizable, and scalable. With accompanying software (https://github.com/jsunn-y/DeCOIL), DeCOIL can be readily implemented to generate desired informed libraries.

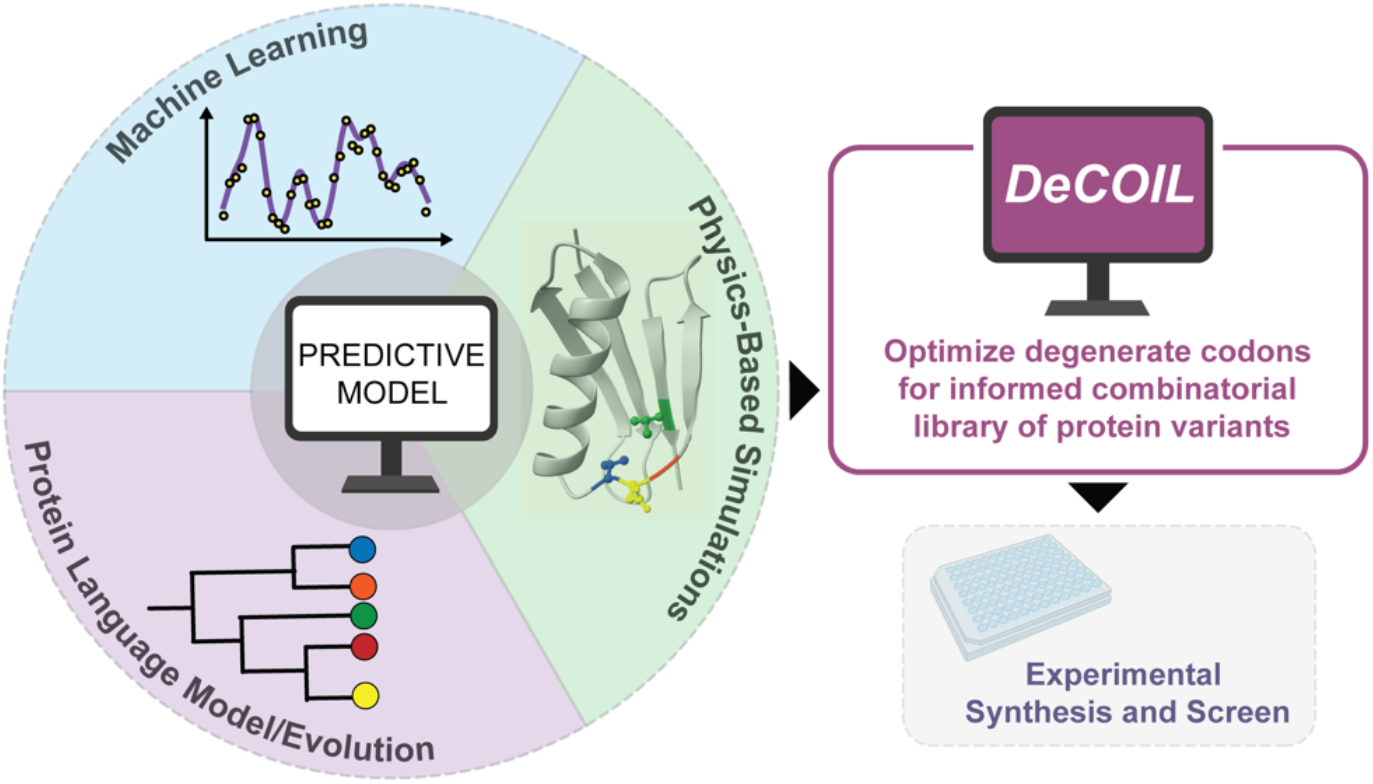

## 1. INTRODUCTION

Proteins can be engineered to improve them for applications ranging from chemical manufacturing to diagnostics and therapeutics.^1–3^ Directed evolution (DE) is a powerful protein engineering method that optimizes protein fitness by greedy hill climbing in amino-acid sequence space. Recently, machine learning (ML) has emerged as a useful tool to complement DE and accelerate protein engineering,^4–13^ as has been done with channelrhodopsins,^14,15^ adeno-associated viruses,^16,17^ enzymes,^18,19^ and other proteins.^5,20,21^ In ML-assisted protein engineering (MLPE), ML models are trained on data to learn a mapping between protein sequences and their associated fitness values to approximate protein fitness landscapes.^22–26^ These trained models can then predict the fitness of previously unseen protein variants, increasing screening efficiency by evaluating proteins *in silico* and expanding exploration to a greater scope of sequences, compared to conventional DE approaches. For instance, in machine-learning assisted directed evolution (MLDE),^27^ an ML model is trained on a small sample of variants in a multi-site simultaneous mutagenesis (combinatorial) library and then used to predict fitness and rank all variants within the combinatorial space. Similarly, “zero-shot” (ZS) scores, such as those based on conservation in evolution or protein-stability calculations, can estimate protein fitness^28–38^ and guide the direction of protein engineering campaigns, without collecting labeled data.^39–41^ These two approaches are combined in focused-training MLDE (ftMLDE), where Wittmann et al. find that sampling training libraries with variants harboring favorable ZS scores yielded ML models with better performance than random sampling.^42^ Overall, with the rise of ML-based and other computational methods, protein engineers are increasingly better informed about what protein variants to explore and screen.

Synthesizing the informed libraries suggested by ML models or ZS scores (predictive models), however, can be non-trivial. The most direct method of variant library construction involves DNA synthesis of each individual sequence, which often remains prohibitively expensive, especially as the number of sequences to be screened increases. Many protein engineering efforts instead utilize DNA library preparation techniques such as degenerate codons (DCs), error-prone polymerase chain reaction (epPCR), DNA shuffling,^43^ or structure-guided recombination.^44^ Early efforts to generate informed protein libraries focused on recombination strategies.^45,46^ Over time, DC, or mixed-base, libraries have been designed manually for DE, using methods such as random “NNK” and the 22-codon trick.^47^ DC libraries are particularly useful for studies targeting mutations to around 4–7 residues simultaneously (combinatorial mutagenesis),^16,27,42^ as in the implementation of MLDE and ftMLDE. **Fig. 1A** highlights steps in possible MLPE workflows where it is desirable to design informed DC libraries (purple arrows). Within these workflows, training libraries are screened and then used to train an ML model, which is used as the predictive model for further round(s) of mutation and screening. By contrast, testing libraries are final libraries, from which an optimized protein variant will be identified. With the expansion of computational models that can predict protein fitness, there is a growing need for computational methods that can optimize DCs for training and testing libraries to sample desirable protein variants suggested by these predictive models.

**Fig. 1.**
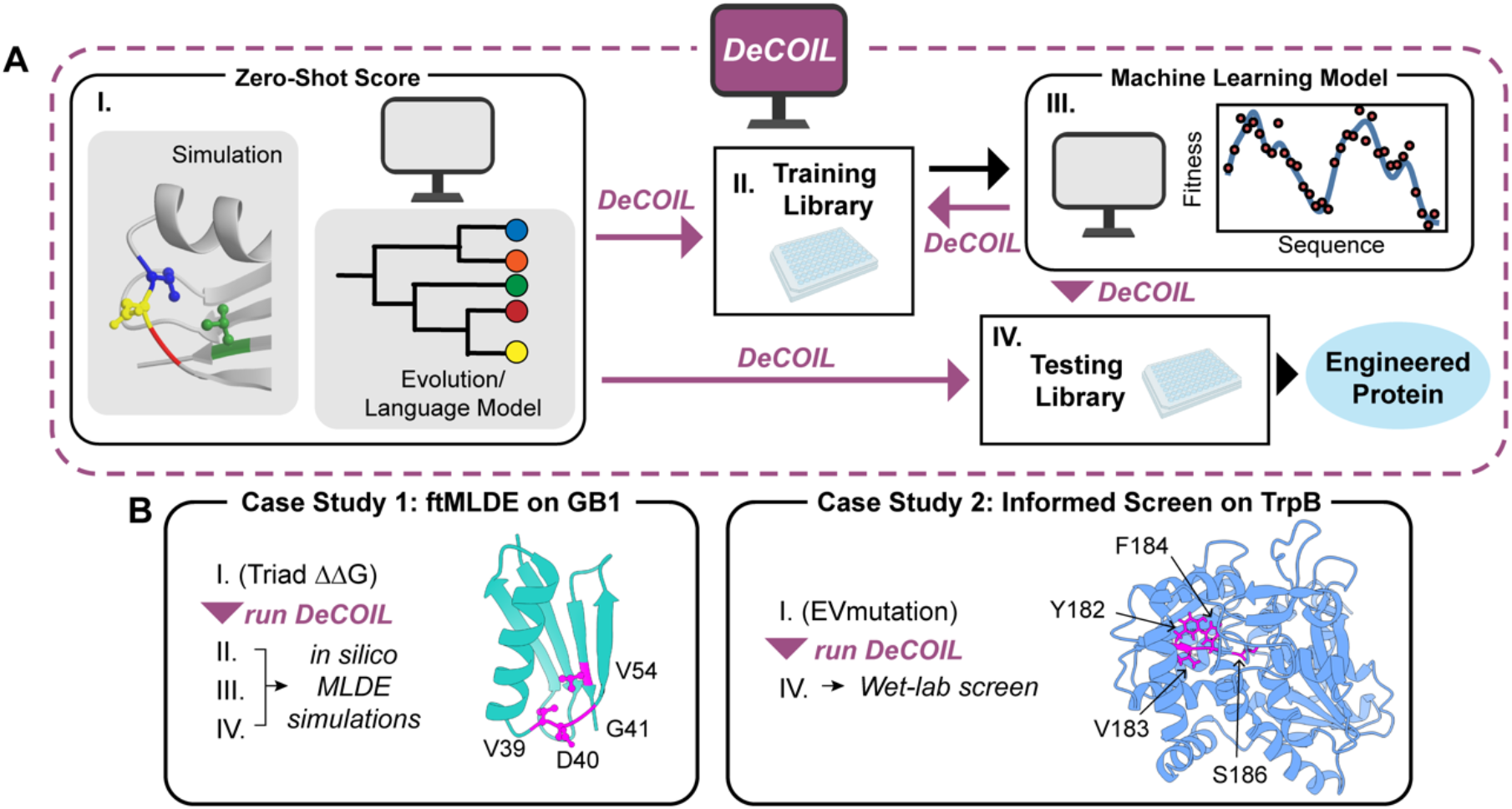
Use cases for DC library design (e.g. DeCOIL) in MLPE workflows. **(A)** DeCOIL can be used to generate a designed library for screening based on computational predictors (ZS scores or ML models) at many possible points along the route to engineering a protein. Examples of workflows: MLDE is II-III-IV; ftMLDE is I-II-III-IV; Bayesian Optimization is II-III-II-III-IV. **(B)** In this study, we explore two case studies. (1) DeCOIL is run between steps I and II on GB1 for designing a training library for ftMLDE based on the Triad ΔΔG of folding between parent and variant protein. Steps II, II, and IV are validated with *in silico* MLDE simulations. (2) DeCOIL is run between steps I and IV on TrpB for designing a testing library based on EVmutation scores. Step IV is validated in the wet lab with a fitness screen. The residues targeted for combinatorial mutagenesis are highlighted in magenta in each protein structure.

Existing methods to design DC libraries, however, are not well suited to utilize the information provided by these predictive models. For example, some DC library optimization approaches (such as LibDesign,^48^ SwiftLib,^49^ and DeCoDe^50^) aim to include as many desired DNA sequences as possible without introducing undesired constructs. However, these approaches are less appropriate for MLPE, where the predictive model does not propose a fixed set of sequences. Other studies take different approaches to bias libraries toward distributions of favorable variants, ranging from theoretical to more applied.^16,51–53^ Each study approaches the problem from a different angle, but none have been made both suitable and practical for the possible use cases encountered during MLPE (**Fig. 1A**).

To address this challenge, we present DEgenerate Codon Optimization for Informed Libraries (DeCOIL). The key contribution of DeCOIL is that it optimizes an objective function directly suited for MLPE–to generate DC libraries that sample sequences biased toward variants likely to have high fitness (exploitation) while maintaining diversity in the sequence search space (exploration). DeCOIL provides several advantages compared to previous methods: (1) it is a general framework that can be applied to many MLPE use cases; (2) the optimization objective explicitly maximizes the value gained by sampling sequences from a DC library; (3) there is consideration of the diversity in types of amino acids covered in the sequence search space; (4) pairwise and higher-order effects (epistasis) between residues is implicitly captured; and (5) the amount of desired exploitation and exploration can be tuned with a single parameter.

In this work, we test DeCOIL on two case studies, as outlined in **Fig. 1B**. We primarily focus on the first, ftMLDE on the B1 domain of protein G (GB1), an immunoglobin binding protein where fitness is measured by binding affinity-based sequence enrichment.^54^ We use DeCOIL to optimize a four-site combinatorial training library to oversample protein variants with favorable ΔΔG values, as calculated by physics-based simulations using Triad side-chain repacking optimization.^55,56^ Each ΔΔG is a computational approximation of the stability of a protein variant, relative to parent. After evaluating with *in silico* simulations, we find that that the optimized DC libraries achieve comparable performance for ftMLDE at a much lower cost than ordering exact sequences. In the second case study, we validate DeCOIL experimentally by synthesizing several DeCOIL-optimized libraries designed to oversample evolutionarily likely variants of the β-subunit of tryptophan synthase (TrpB) from *Thermotoga maritima*, an enzyme that catalyzes the formation of tryptophan from indole and L-serine.^57,58^ Because tryptophan synthesis is the native activity of TrpB, we use the EVmutation Potts model^28^ as a predictive model for protein variant fitness. Through wet lab experiments, we show that the four-site simultaneous mutagenesis libraries are significantly enriched in variants with higher native enzymatic activity, compared to random (NNK) libraries. DeCOIL scales well computationally and generalizes to many applications. Code is readily available at https://github.com/jsunn-y/DeCOIL with a step-by-step user guide. Accompanied by cost-effective sequencing of every variant in a protein library (evSeq),^59^ DeCOIL unlocks a key component for the practical end-to-end implementation of many MLPE workflows.

## 2. RESULTS & DISCUSSION

### 2.1 DeCOIL for optimizing DC libraries

DeCOIL outputs optimized DC libraries, where each library is designed to maximize the average utility of sets of sequences sampled from that library. A DC library is defined by one or more DC templates, where an example of a single DC template for a four-site combinatorial library is NNK-NNK-NNK-NNK. In this study, unless otherwise specified, a DC library refers to a single DC template. Utility for MLPE can be thought of as a form of “weighted coverage” which balances high fitness and diversity (**Materials and Methods**). *Coverage* describes how well a DC library covers the combinatorial protein sequence search space under exploration, and each sequence to be covered is *weighted* by the value from a predictive model such as a ZS score or an ML model (higher weights are given to sequences favored by the model). Having high coverage is desirable for a library so that the sequences sampled are diverse (exploration), and having variants with high weights means they will be more likely to have high fitness (exploitation). Thus, the optimization objective for DeCOIL is both intuitive and well suited for MLPE by balancing exploration and exploitation.

A single parameter, *p*, can be used to tune the desired level of exploration and exploitation, when running DeCOIL (**Fig. 2**). In this work, *weight* is defined as the normalized ranking of a variant from the predictive model raised to the power *p* (**Materials and Methods**). As *p* is increased, the relative value of the most favorable variants from the predictive model will be increased, thus a smaller region of the search space will be considered valuable. A small value of *p* implies a focus on exploration, or a desire for broader coverage, while a large value of *p* implies a focus on exploitation, which signifies more trust in the predictive model.

**Fig. 2.**
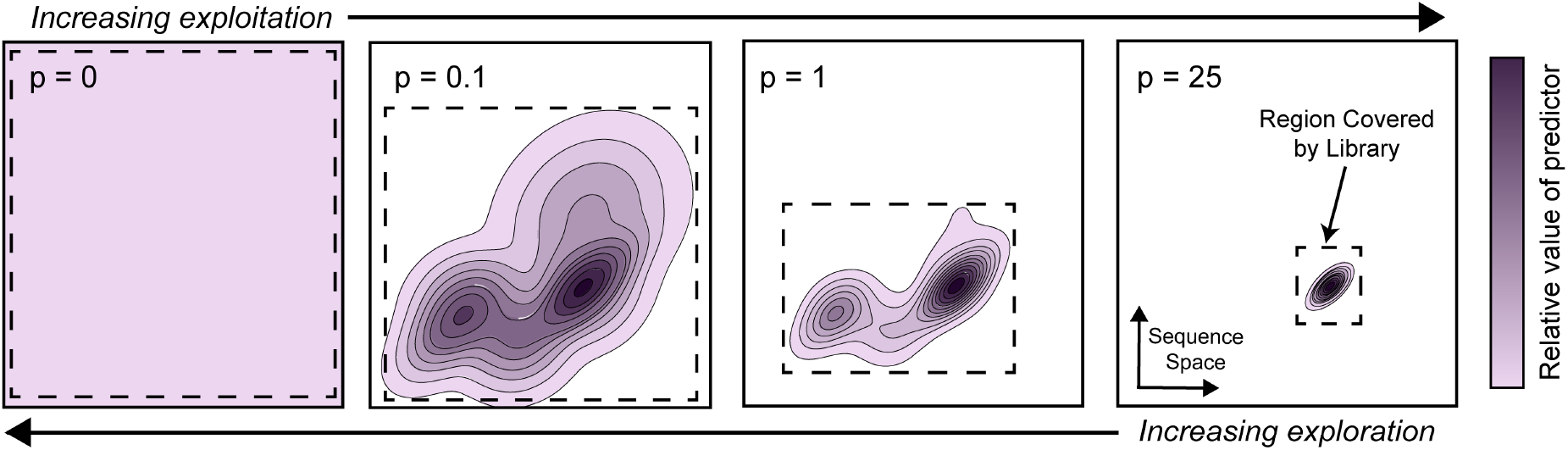
A single parameter, *p*, controls the balance of exploration and exploitation for library optimization using DeCOIL. DeCOIL optimizes an objective that aims to cover sequences that are deemed to have high value by the predictor (predictive model). As illuminated by a hypothetical predictor in purple, increasing *p* will increase the relative value of the most favorable variants from the predictor, constraining the diversity of desirable sequences. In other words, increasing *p* increases exploitation while decreasing exploration. If *p* = 0, then the objective is unweighted (unbiased exploration).

When designing a library, it is important to balance exploration differently for different predictive models (**Fig. 3)**, as an ideal library will lie somewhere between these the two extremes of one that is fully explorative (such as NNK) and one that is fully exploitive. For a predictive model that is weakly correlated to the true protein fitness landscape, exploration is preferred; otherwise, the fitness landscape cannot be learned, and the optimum may be missed. Exploration is also beneficial when constructing a training library, as the model can better interpolate within its broader training feature space, rather than extrapolating to unseen sequences. For a predictive model that is strongly correlated to the true protein fitness landscape, such as an ML model trained on sequence-fitness pairs, exploitation will be preferred to best enrich the proposed library in high-fitness candidates. In this case, too much exploration will result in a library with many low-fitness variants, wasting resources and increasing the required screening burden. Thus, exploitation may be preferred for a testing library. DeCOIL users may not know the ideal value of *p* to use for their given exploration-exploitation tradeoff, but they can make an informed decision considering the reasoning above.

**Fig. 3.**
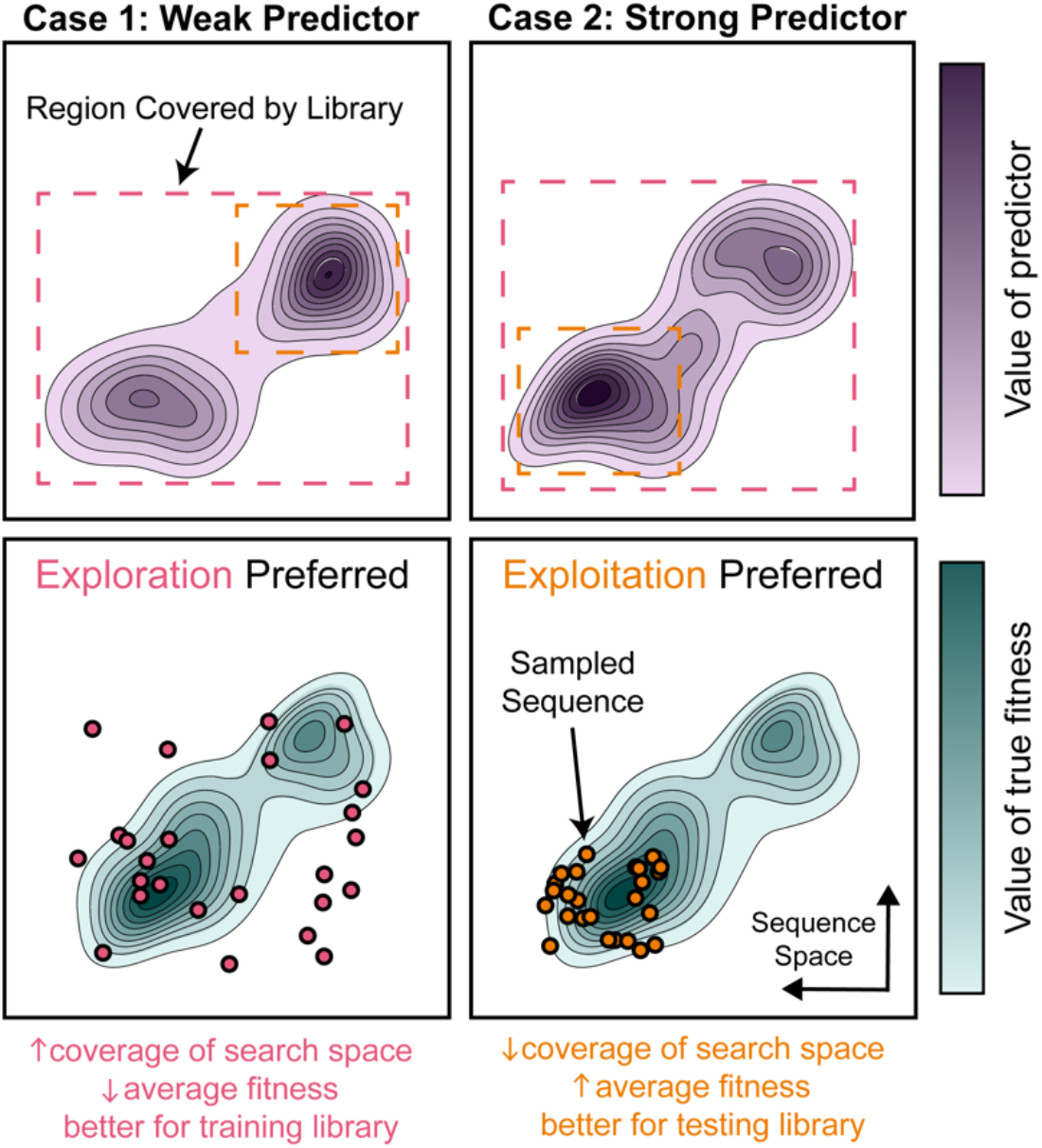
Hypothetical landscapes demonstrating where exploration or exploitation is preferred when designing a protein variant library. In the first column, exploration (magenta) is preferred because the predictor is weakly correlated to the true fitness distribution. In the second column, exploitation (orange) is preferred because the predictor is strongly correlated to the true fitness distribution.

DeCOIL optimizes each DC library using greedy hill climbing; in each iteration a small change is made to which nucleotides are allowed in the library, and the change is only accepted if the set of sampled sequences has higher weighted coverage (the objective) on average than the previous library (**Materials and Methods)**. During optimization, to decrease computational cost, our optimization objective uses a simplified notion of coverage termed “step” *coverage*, which is directly proportional to the number of unique sequences sampled by the library. However, during evaluation of optimized libraries, we additionally use “diffuse” *coverage*, which adds additional complexity to account for amino acid types (**Materials and Methods**). A visual explanation of the difference between step and diffuse coverage can be found in **Fig. S2** of **Supporting Information**.

DeCOIL scores based on sampling the target number of variants to be screened, so it directly captures how much utility is gained from screening a library of a specified size. Thus, DeCOIL implicitly accounts for pairwise and higher order covariation in amino acid substitution effects– when mutation effects at different residues are not independent (epistasis). A hypothetical example of this effect is given in **Fig. S1** of **Supporting Information**. The optimization configuration is generalizable to (1) different experimental library screening sizes and (2) any predictive model that assigns a relative value to each protein variant in the search space. Finally, the computational cost of DeCOIL scales linearly with the number of sites in the combinatorial library, making it computationally efficient. Together, these qualities make DeCOIL appropriate for a variety of design tasks.

### 2.2 Analysis of optimized DC libraries on GB1

The next two sections will focus on the first case study: using DeCOIL to optimize training libraries for ftMLDE on a four-site combinatorial mutagenesis landscape determined for GB1.^54^ **Table 1** summarizes the average performance of optimization solutions from three DeCOIL campaigns, each using a different value of *p*. The DeCOIL-optimized libraries perform significantly better than a random NNK library across several metrics, including weighted step coverage and the number of unique sequences in the library. The optimized libraries generally only sample a small fraction of the total amino acid sequence search space (**Fig. S3, Supporting Information**). The optimized libraries are also significantly enriched in variants that are within the top 95^th^ percentile of Triad ΔΔG scores, and because the Triad ΔΔG is correlated to binding fitness, the optimized libraries sample variants with generally higher fitness. Tuning the *p* value also demonstrates the exploration-exploitation tradeoff. As *p* is increased, the fraction of unique sequences in the library decreases, but the fraction of sequences in the top 95^th^ percentile of Triad scores increases, as does the average fitness of variants in the library. Impressively, DeCOIL can propose libraries where over 71% of the screened variants are unique and in the top 95^th^ percentile of Triad ΔΔG scores. We further show that DeCOIL proposes better solutions than those from the DC optimization software Swiftlib,^49^ under somewhat equivalent specifications and parameters (**Table S1, Supporting Information**). Optimization allowing more than one DC template per library yields designs that perform slightly better, with diminishing returns as additional templates are used (**Table S1, Supporting Information**). We also attempted to compare our method to DeCoDe,^50^ but that algorithm did not converge to a solution given the targets provided. Other relevant methods were not available to us for comparison at the time of this study.^16^

**Table 1.**
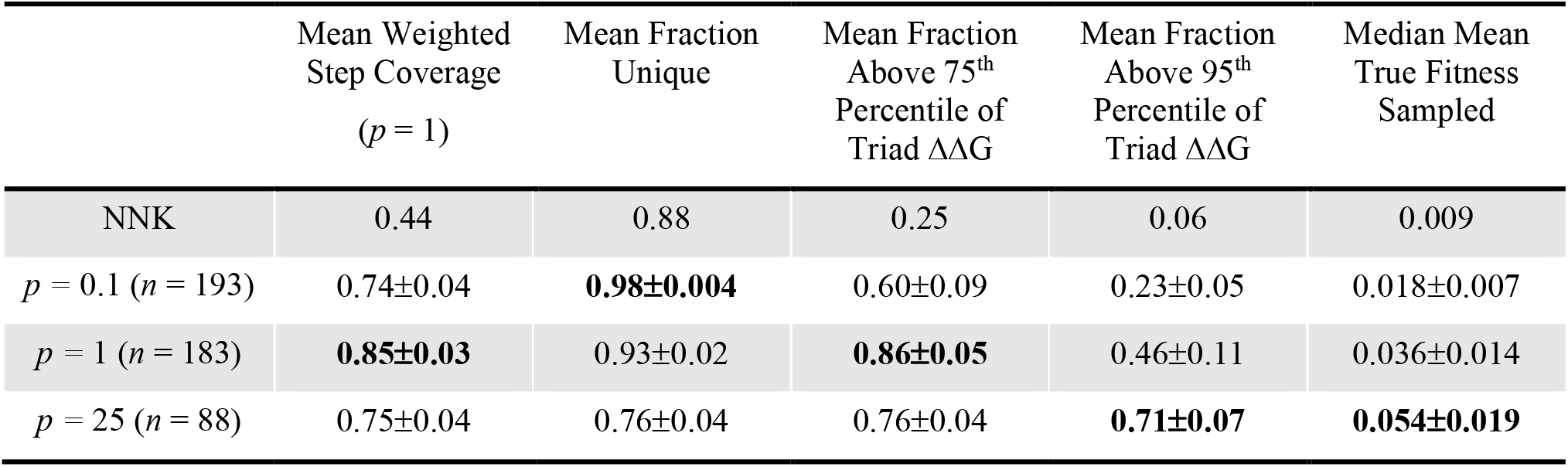
Performance of optimized DC libraries, on GB1. Optimized libraries using different values of *p* are skewed toward desired protein distributions, as measured by several metrics. The weighted step coverage of the libraries, calculated with *p*=1, is given. As *p* is increased, the fraction of unique sequences in the library decreases, but the fraction of unique sequences in the top 95^th^ percentile of Triad ΔΔG scores increases, as does the median mean fitness of the library (not used during DeCOIL optimization). *n* refers to the number of unique solutions (based on amino acid distribution) after optimization from 240 random initializations. Mean fraction refers to the fraction of the screening library that is unique sequences, for sequences not containing stop codons. All optimized libraries use the Triad ΔΔG rank on the GB1 dataset with a screening size of 384.

The greedy hill climbing optimization trajectories from the libraries summarized in **Table 1** are presented in **Fig. 4A–B**, where we see that the lower the *p* value, the faster the average convergence to a solution. This suggests that libraries with lower *p* values are easier to optimize and more similar to random initializations. **Fig. 4C** shows the average Triad ΔΔG rank distribution of libraries for each value of *p*, and **Fig. 4D** shows each of their average true fitness rank distributions. The libraries optimized using larger *p* values are more skewed toward variants with the most favorable Triad scores, and by extension the variants with highest fitness.

**Fig. 4.**
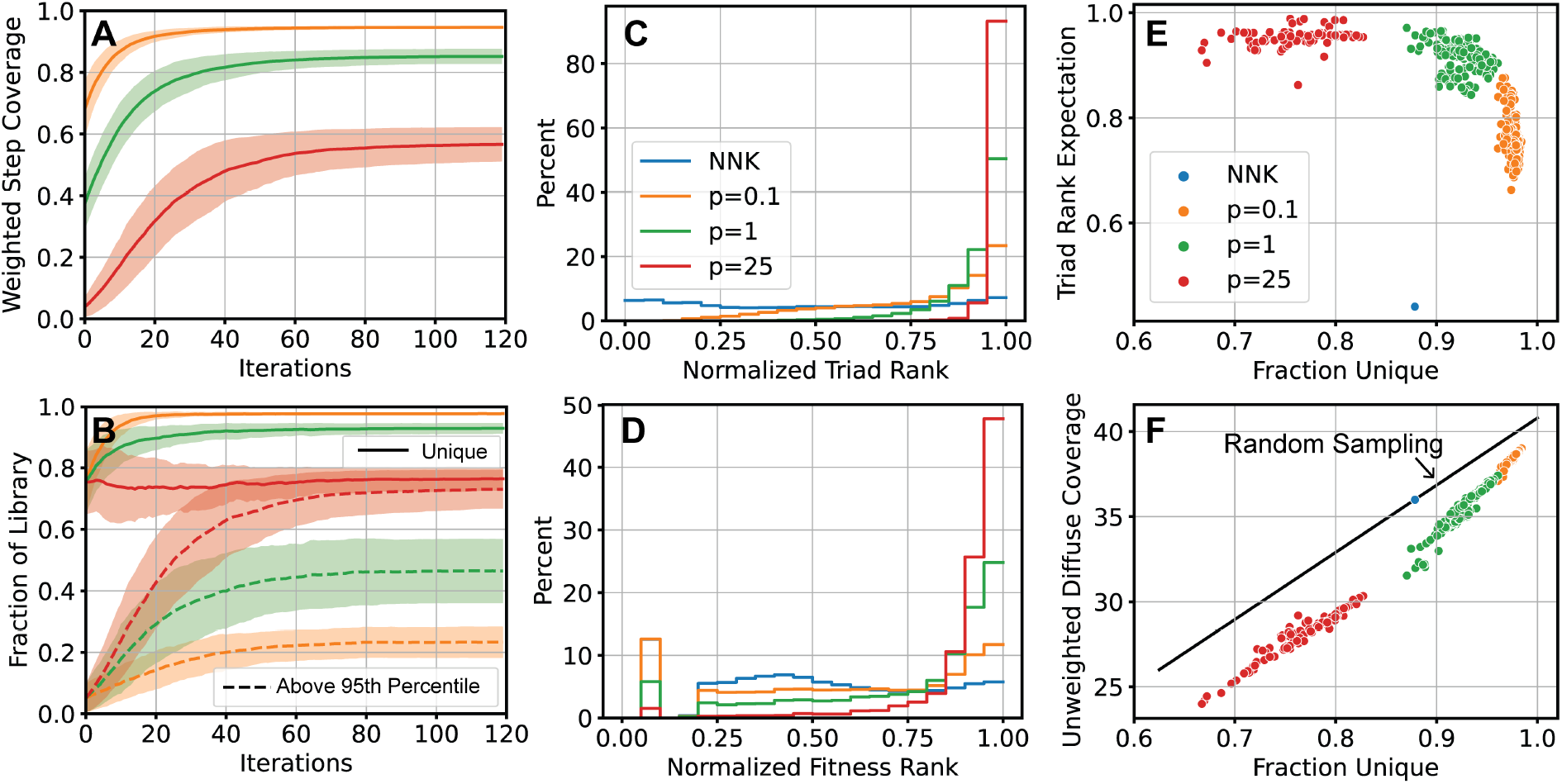
Optimized DC libraries show enhanced fitness and high sequence-space coverage, but inherent tradeoffs, on the GB1 landscape. Optimization trajectories of DC libraries, showing **(A)** weighted step coverage (the optimization objective) with different values of *p* and **(B)** the fraction of sampled libraries that are unique protein variants (solid) and that are unique variants in the top 95^th^ percentile of Triad ΔΔG scores (dashed). Histograms showing that average optimized libraries are significantly enriched in variants with **(C)** favorable Triad rankings and **(D)** fitness rankings (unseen during training), compared to the random NNK library. **(E)** Tradeoff between exploitation (y axis) and exploration (x axis): for a given library, the expectation of the normalized Triad rank decreases as the number of unique samples in the library is increased. Libraries optimized using *p* = 25 are expected to have higher Triad scores, but lower coverage of the search space. Libraries optimized using *p* = 0.1 are expected to have higher coverage of the search space, but lower Triad scores. **(F)** For randomly sampled exact libraries, unweighted diffuse coverage increases linearly as a function of the number of unique samples (black line), with the NNK library lying on this line. Optimized DC libraries have lower diffuse coverage of sequence space than random sampling (for a given fraction of unique variants in the library), which is an inherent limitation of using DCs, as they constrict residues to certain amino acids. Diffuse coverage is calculated using Hamming distance and σ=0.4. All optimized libraries use the Triad ΔΔG rank on the GB1 dataset with a screening size of 384. Fraction unique is directly proportional to unweighted step coverage.

The optimized libraries also demonstrate inherent tradeoffs. As shown in **Fig. 4E**, as the fraction of unique sequences in the library (exploration) increases, the expected normalized Triad rank to be sampled from the library (exploitation) decreases. Libraries optimized with small *p* values thus favor exploration at the expense of exploitation, while the opposite is true for libraries optimized with large *p* values. Additionally, **Fig. 4F** shows the loss of sequence space coverage that occurs when using DC libraries. Unweighted diffuse coverage, calculated using Hamming distance and σ=0.4, increases linearly as the fraction of unique sequences in a library for randomly sampled libraries (black line). NNK libraries fall on this line. However, all optimized DC libraries lie below this line because, at each residue, a DC library is constrained to discrete amino acid possibilities, which reduces diffuse sequence-space coverage. This inherent penalty suggests that exact sequences will always offer at least some advantage in terms of sequence-space coverage, compared to DCs.

Unlike many existing DC-optimization algorithms, which are largely deterministic, DeCOIL can find many solutions to the same problem that are quite different from each other, though they share common learned patterns (**Fig. S4, Supporting Information**). This is particularly useful because protein engineers can visualize these solutions, rather than choosing one blindly. Furthermore, the designed libraries can be assessed by calculating other evaluation metrics or infusing domain knowledge before selecting the library to use for an experimental screen. In the next sections, we explain how we employ this to choose optimized DC libraries with ideal downstream performance.

### 2.3 Optimized DC libraries perform better for downstream MLDE on GB1

After optimizing hundreds of DC libraries in parallel, we evaluated their downstream performance by simulating 24 ftMLDE campaigns *in silico* for each library. The violin plots in **Fig. 5A–B** show the performances of DC libraries, given by the median of both the max and mean fitness achieved from sampling training sets of 384 sequences from the libraries, training ML models on each of these sets, and evaluating the top 96 predictions from each of the proposed test libraries. On average, the libraries optimized using *p* = 0.1 or *p* = 1 perform better than random and better than the top designs from Swiftlib proposed under somewhat equivalent specifications and parameters (**Fig. S5, Supporting Information**). The DeCOIL library designs using multiple templates offer slightly improved performance, on average (**Fig. S6, Supporting Information**). However, the libraries optimized using *p* = 25 perform quite poorly for ftMLDE despite their clear bias for the best Triad scores and fitness values. The low performance of ML models trained on libraries optimized with *p* = 25 suggests that these libraries have poor coverage of the search space, overfitting to the Triad scores and sampling from constrained regions of the search space that miss the highest fitness variants. Thus, for training library construction with weak predictors, we propose prioritizing exploration, a claim which is further supported by the results in **Fig. S7** of **Supporting Information**. A similar phenomenon is observed in previous work.^60^

**Fig. 5.**
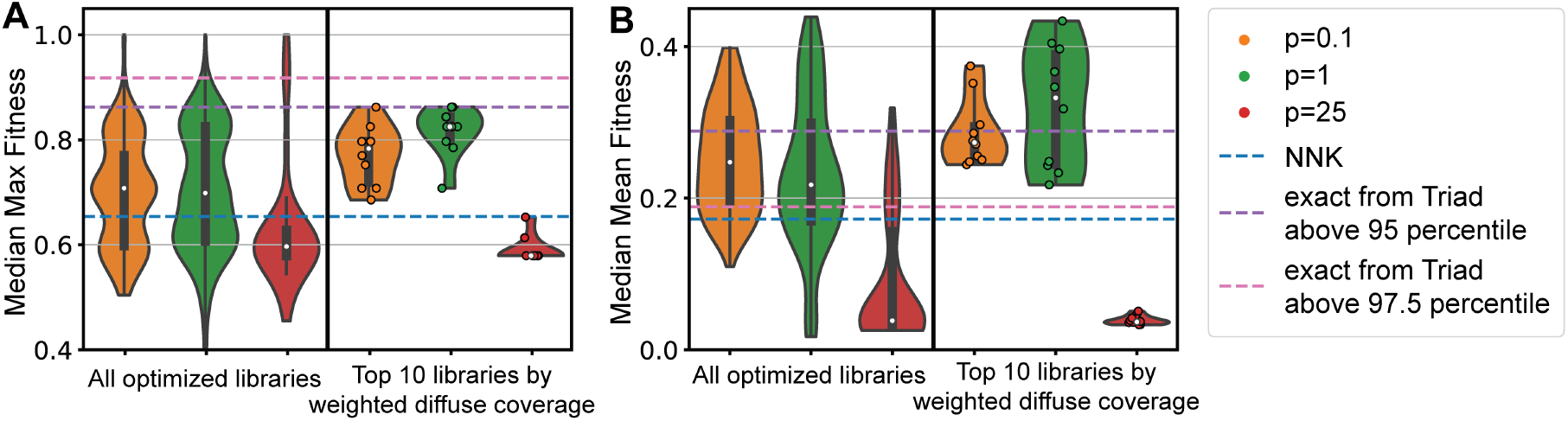
Computational simulations show that optimized libraries are generally more effective than random sampling for MLDE, on GB1. Median **(A)** max and **(B)** mean fitness achieved from downstream MLDE simulations, using proteins sampled from optimized DC libraries as training sets. The violins in the first column correspond to all of the optimized libraries, and those optimized using p=0.1 or p=1 perform better than random (NNK). Selecting the top 10 libraries for the p=0.1 or p=1 optimization campaigns, based on weighted diffuse coverage, boosts the performance significantly to where it is comparable to the original demonstration of ftMLDE using exact sequences;^42^ these are provided as dashed baselines for randomly sampled exact training sets from the top 95 and 97.5 percentile of Triad scores. Diffuse coverage is calculated using Hamming distance and σ=0.4. Each ML model is an ensemble of five boosting models; training sets of 384 sequences (not all unique) were sampled from different libraries with 24 repeats; max and mean fitness were evaluated from the top 96 predictions of each ML model and normalized to the maximum fitness in the landscape. Tests were performed on the GB1 dataset, using the Triad ΔΔG rank as a ZS score.

While the DC library solutions with *p* = 0.1 or *p* = 1 perform better than random libraries on average, we reason that those with more favorable mean Triad ΔΔG scores and greater diversity would be more effective. To test this hypothesis, we measured the correlation between various metrics and downstream MLDE performance, for the optimized libraries. In **Fig. S7** of **Supporting Information**, we show that MLDE performance, measured by median max and mean fitness achieved, is most strongly predicted by weighted diffuse coverage, calculated using Hamming distance and σ=0.4. This conclusion supports our claim that weighted diffuse coverage is a strong metric for utility in MLPE. Weighted diffuse coverage considers exploitation of a predictive model and exploration of the search space with consideration of amino acid types, and it is more correlated with MLDE performance than other more simplified metrics. Therefore, we selected the top ten unique optimized DC libraries from each optimization campaign with *p* = 0.1 or *p* = 1 using weighted diffuse coverage based on Hamming distance and σ=0.4. These selected libraries indeed demonstrate significantly higher average performance for MLDE, as shown in the second column of **Fig. 5A-B**. In fact, the performances of these selected libraries approach or exceed those of libraries generated from exact sequences using percentile cutoffs based on Triad ΔΔG score (**Fig. S5, Supporting Information**), which was the previous standard which led to ideal performance for ftMLDE.^42^ However, it is not entirely obvious why certain DeCOIL-optimized libraries perform better for MLDE (**Fig. S8, Supporting Information**).

Ultimately, DeCOIL enables protein engineering using ftMLDE with comparable outcomes, without the need for ordering costly exact sequences for the training library. We would also like to note that the simulated performance of these libraries is likely an underestimate because approximately seven percent of the full landscape lacks fitness labels. In the case that one of these sequences is missing a label, it is dropped from the training set, reducing the number of sequence-fitness pairs used for model training.

### 2.4 Experimental screens of optimized DC libraries on TrpB

In the second case study, we used DeCOIL to optimize testing libraries based on ZS scores predicted by EVmutation, on TrpB. We then performed experimental validation by synthesizing four-site simultaneous site-saturation libraries, sampling variants from them, and measuring their associated rates of tryptophan formation (fitness). We performed two DeCOIL campaigns with *p* = 1 and *p* = 25, each starting from 240 random initializations. The objective, weighted step coverage, was maximized over 120 iterations of greedy climbing, with the *p* = 1 campaign converging somewhat more rapidly than the *p* = 25 campaign (**Fig. 6A**). During these campaigns, the fraction of unique protein variants sampled by the library increased, as did the fraction of variants above the 95^th^ percent of EVmutation scores (**Fig. 6B**). The different optimized libraries are visualized in **Fig. S9** of **Supporting Information**. We selected three optimized libraries, with their corresponding degenerate codons and amino acid distributions shown in **Fig. 6C**. One library was selected from the DeCOIL campaign with *p* = 1 (Lib1), while two libraries were selected from the campaign with *p* = 25 (Lib2 and Lib3). We chose these libraries to explore further because the three libraries resemble each other, but Lib1 encouraged more exploration at residues 182 and 186 compared to Lib2 and Lib3. Furthermore, some of our unpublished experiments with TrpB suggest that residues 182 and 186 are generally restricted to a smaller subset of amino acid types to preserve function. The expected distribution of EVmutation scores of these selected libraries is shown in **Fig. 6D**. Lib2 and Lib3 (from *p* = 25) are more biased toward variants with the most favorable EVmutation scores compared to Lib1 (from *p* = 1). The random NNK library displays a very slight bias toward evolutionarily favorable variants.

**Fig. 6.**
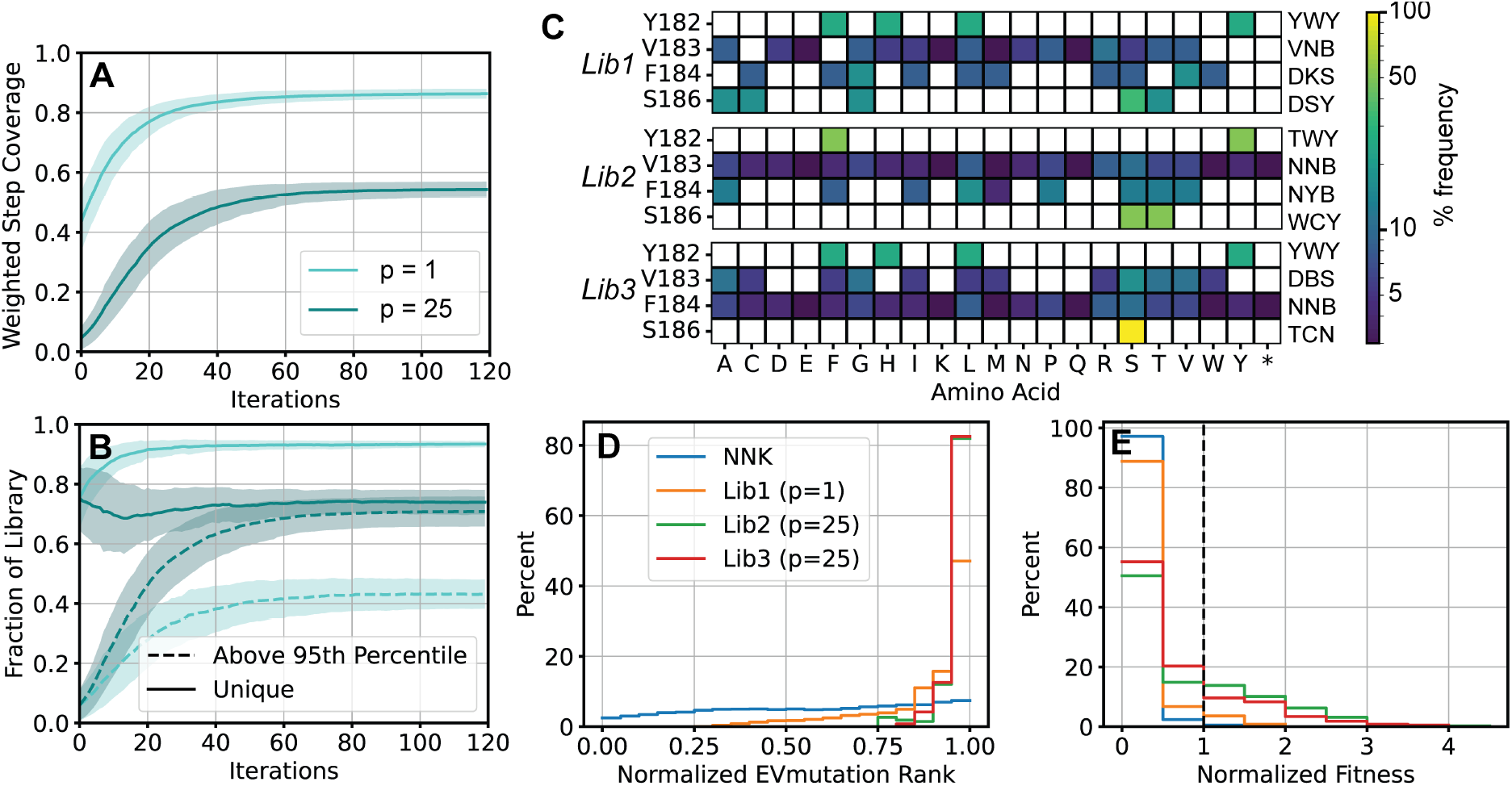
Performance of optimized libraries on TrpB. Optimization trajectories of DC libraries, showing **(A)** weighted step coverage (the optimization objective) with different values of *p* and **(B)** the fraction of sampled libraries that are unique protein variants (solid) and that are unique variants in the top 95^th^ percentile of EVmutation scores (dashed). **(C)** Three selected libraries, defined by IUPAC nomenclature, and their corresponding amino acid distributions. Lib1 comes from the optimization campaign with *p* = 1, and Lib2 and Lib3 come from the optimization with *p* = 25. Histograms showing that selected optimized libraries are significantly enriched in variants with **(D)** favorable EVmutation rankings and **(E)** fitness values, compared to the random NNK library. Fitness is normalized to the parent proteins on each plate. Tests were performed on TrpB, using the EVmutation rank as a ZS score and fitness as the rate of tryptophan formation. An asterisk is used to denote stop codons.

To validate the optimized libraries, we performed experimental screens on sets of 384 variants sampled from each library. The fitness distributions of the sets of variants are skewed toward higher fitness (**Fig. 6E**). Lib1 has a small fraction of variants with activity, and a few of these achieve better fitness than parent. Lib2 and Lib3 are significantly enriched in high fitness variants, with only around 50% of the sampled variants close to inactive. By contrast, the NNK library is nearly entirely comprised of inactive variants (zero-fitness), and none of the sampled variants have fitness higher than parent. The libraries from optimization campaigns with *p*=25 (Lib2 and Lib3) identified more higher fitness variants than those from *p*=1 (Lib1), so we argue that exploitation is preferred for this use case. Exploitation is likely preferred because we are (1) designing a testing library, and (2) the predictive model (EVmutation) is strongly correlated to fitness (native tryptophan formation). Impressively, the libraries proposed by DeCOIL achieved significant gains to fitness over parent, despite only screening 384/160,000 = 0.2% of the search space, which contains mostly dead variants. Overall, these results suggest that DeCOIL can enable protein engineering discovery by helping with the design of testing libraries enriched with higher fitness variants.

## 3. CONCLUSION

DeCOIL is a computational method to optimize DCs for synthesizing an informed combinatorial mutagenesis protein variant library, which is a critical step in a variety of MLPE workflows. DeCOIL aims to bias a DC library toward desired proteins based on a predictive model from physics-based simulations, evolutionary knowledge, or ML–while sacrificing minimal coverage of the sequence search space under exploration. Thus, DeCOIL directly optimizes the utility of screening a sample of sequences from a library for MLPE. DeCOIL can play a role in informed library screening and many MLPE workflows, such as MLDE and ftMLDE. It can be used to generate training libraries as inputs to ML models, or testing libraries where the goal is simply to identify the protein variant with the highest fitness. At the same time, DeCOIL is simple; with a single parameter, it can be tuned to favor exploration for training libraries and exploitation for testing libraries based on increasing trust in the predictive model.

We establish that DeCOIL is useful for both training and testing libraries. DeCOIL can skew a training library to achieve higher fitness for ftMLDE without the need for ordering costly exact DNA sequences, as we demonstrate by optimizing using the Triad ΔΔG score and validating using *in silico* MLDE simulations on the GB1 landscape. On the other hand, DeCOIL can significantly enrich a testing library in higher fitness variants, which we demonstrate by optimizing TrpB libraries to oversample variants with high EVmutation scores and experimentally validating the enzymatic activity of variants sampled from those libraries.

DeCOIL is generalizable to any predictive model and library screening size, all while accounting for pairwise and higher order amino acid substitution effects. Furthermore, greedy hill climbing optimization is efficient and scales linearly with the number of residues in the combinatorial library and the number of DC templates desired in a library. Future work could involve generalizing DeCOIL to other DNA synthesis technologies. We expect that DeCOIL, combined with sequencing of every variant in a protein library (evSeq), will be useful in a variety of MLPE workflows.

## 4. MATERIALS AND METHODS

### 4.1 Utility for MLPE as the library optimization objective

We present a generalized optimization objective that is directly aligned with the goals of library design for MLPE. When sampling a set of amino acid sequences 𝒜 from an input library *L*, we desire that this set of sequences: (1) has high average fitness based on a predictive prior and (2) has high diversity in sequence space (covers the input search space well). This objective can be mathematically formulated as maximizing the objective function *J* (overall weighted coverage), which is defined in **Eq. 1** as the sum across all sequences *i* in the total search space of sequences 𝒰 of a predictive-score-dependent weight (*w*_*i*_) multiplied by the set coverage (*C*_*𝒜*_(*i*)) of sequence *i* by the set of sequences of *𝒜*.

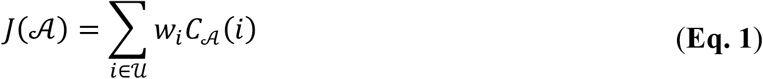

𝒰 can typically be thought of as all sequences in the combinatorial mutagenesis space of sites under exploration, or as a relevant subset of this total search space. *w*_*i*_ = *f*(*Z*(*i*)) In other words, each weight *w*_i_ of sequence *i* should be a function *f* of its sequence’s predictive score (*Z*(*i*)). In this work, we typically use a weighted coverage where we treat *f* to be the normalized ranking of *Z*(*i*), raised to the power *p*. Thus, the variant with the lowest (least favorable) predictive score is given *w* = (1/*N*)^*p*^ and that with the most favorable ranking is assigned *w* = (*N*/*N*)^*p*^ = 1, where *N* is the total number of sequences in 𝒰. If *p* = 0, *f*(*x*) = 1, then the coverage will be unweighted. A visualization of the weight function is provided in **Fig. S10A** of **Supporting Information**.

The set coverage *C*_*𝒜*_(*i*) of sequence *i* by the set of sequences of 𝒜 can be further defined, as in **Eq. 2**.

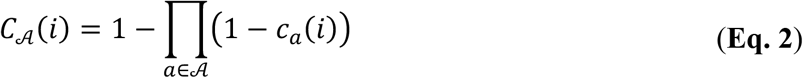

This is the complement of the product of the individual coverage, *c*_*a*_(*i*), complements of every sequence *a* in *𝒜*. The individual coverage *c*_*a*_(*i*) is a measure of how similar sequence *a* is to sequence *i*; in other words, it is a measure of how well *a* covers *i*. In this work, we primarily explore two different individual coverage functions:

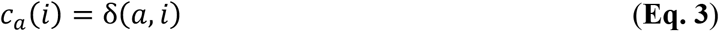

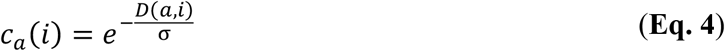

**Eq. 3**, the delta function, is referred to as step coverage, where *c*_*a*_(*i*) = 1 if *a* = *i* and *c*_*a*_(*i*) = 0 otherwise. The second coverage is referred to as diffuse coverage, where individual coverage is an exponentially decaying function of the distance between *a* and *i* (**Eq. 4**). In this case, the hamming distance between amino acid sequences is an appropriate distance function *D*. The Manhattan or Euclidean distance can also be used for distances between continuous embeddings of sequences. σ can be used to tune the width of the tail in this decaying exponential. A visualization of the individual coverage functions used in this work is provided in **Fig. S10B** of **Supporting Information**.

Putting it all together, the optimization objective (**Eq. 5**) is to find the DC library *L*^*^ that satisfies the following:

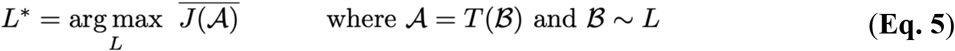

In other words, we are trying find the DC library that has the largest average value (across *n*_*repeats*_) of the objective function *J* across sets of nucleotide bases ℬ containing *n*_*samples*_ sampled from *L* and translated (*T*) to sets of amino acid sequences *𝒜*.

*L* can be composed of one or more templates; NNK-NNK-NNK-NNK is an example of a template for a four-site amino acid combinatorial library. If a library contains more than one template, the total set of sequences sampled will be split evenly such that the same number of sequences is sampled from each template. Because this scoring objective uses sampling, it implicitly accounts for pairwise effects and higher order covariation, which is an advantage. This type of objective is related to submodular set functions.^61^ Submodularity can be intuitively explained by the fact that a new sequence, *s*, will provide more coverage of sequence space if it is added to a smaller existing set, compared to a larger set of sequences. Though maximizing submodular functions is NP hard,^62^ it has been shown that applying a greedy algorithm will provide a close approximation of the optimal value.^63^

### 4.2 Greedy hill climbing optimization

In the case of a single template per library, the input DC libraries are initialized as *n* = 240 random templates. Each template is represented as an encoding of length 12 × k bits, where k is the number of the amino acid sites in the combinatorial library. Each 12-bit representation is a degenerate codon, or a triplet of 4-bit representations, each representing if nucleotides A, C, G, and T are allowed, respectively. In each iteration of optimization, *d* = 4 random bits are flipped, and each of these new libraries is evaluated based on the scoring function described in the previous section. Both use cases explored here involve four-site combinatorial mutagenesis libraries, with a screening size of *n*_*samples*_ = 384 and repeated *n*_*repeats*_ = 100 times to reduce uncertainty in the scoring function. The library with the best score is then used as the starting point for the next iteration. The optimization performed uses weighted step coverage as the objective, as diffuse coverage calculations are too computationally demanding on most systems. For the GB1 case study, the ZS score used is the ΔΔG of folding between parent and each variant as calculated using the Triad package from Protabit using fixed-backbone side chain repacking.^55,56^ Three different optimization campaigns are executed, with *p* = 0.1, *p* = 1, and *p* = 25. For the TrpB case study, the ZS score used is the EVMutation score of each variant calculated from a Potts model trained on the multiple sequence alignment of the parent protein,^28^ and *p* = 1 and *p* = 25 are explored. The alignment is generated from the EVcouplings webapp. Optimization is performed for 120 iterations.

DeCOIL also supports optimization of DC libraries with multiple templates per library. In each iteration of optimization, *d* = 4 random bits are flipped in each template. The 384 samples used for scoring are then equally split across each of the templates, for simplicity. Thus, the computational cost of the method scales linearly with the number of sites in the combinatorial mutagenesis library and the number of templates desired per library.

### 4.3 MLDE simulations on GB1

The empirical fitness landscape for GB1 is used, where variation occurs across four epistatic residues and fitness is measured as binding affinity by enrichment of protein bound to the IgG-Fc antibody, using mRNA display coupled to next-generation sequencing.^54^ For consistency, all ML models used in the simulation of MLDE on the GB1 landscape have the same architecture. For each DC library, 24 sets of 384 sequences are sampled from the library and mapped to unique and labelled sequences. Note that the GB1 dataset is only about 93% complete (149,361/160,000 possible sequences have labels), so some of the unique sequences will be dropped during training. Each of the 24 models is an ensemble of five gradient boosting models using XGBoost^64^ with a 90:10 split for training versus validation (for early stopping). The inputs to the model are one-hot amino acid encodings.

### 4.4 Cloning and expression of TrpB enzymes

To prepare the plasmid libraries, we begin with the parent enzyme in a pET22b(+) backbone. From this plasmid, we amplify two individual fragments (fragments A and B) with a break in the ampicillin resistance gene and a gap where the variation will be introduced to reduce parent bias. Fragment A is then used in a second amplification with degenerate primers encoding variation at positions 182, 183, 184, and 186 of the TrpB gene; this creates fragment C, which now consists of a library of sequences. Fragments B and C are then assembled using the Gibson assembly method.^65^ The PCR protocols and primers design are detailed in **Table S2-3** of **Supporting Information**.

The Gibson products are transformed into T7 express competent *Escherichia coli* cells (NEB C2566). Individual colonies are grown in 600 μL Luria Broth (RPI L24400) supplemented with 100 μg/mL carbenicillin (AG Scientific C-1385) in 96-well polypropylene plates overnight at 37 °C, 220 rpm, 80% humidity. Twenty μL of starter culture is used to inoculate 630 μL of Terrific Broth (RPI T15000) supplemented with 100 μg/mL carbenicillin. Cultures are grown for 4 hours at 37 °C, 220 rpm, 80% humidity, then chilled on ice for 15 minutes before induction. Protein expression is induced with the addition of 50 μL of IPTG (GoldBio 367-93-1) to a 1 mM final concentration. Protein expression is carried out at 22 °C, 220 rpm for 20–24 hours. The cells are collected by centrifugation at 4,500 *g*, 4 °C for 10 minutes. The cell pellets are frozen in the 96-well plates at -20 °C until further use, or for at least 16 hours to allow for complete freezing.

### 4.5 Screening of TrpB libraries

TrpB pellets in 96-well plates are thawed in a room temperature water bath for 1 hour. They are next lysed in 300 μL lysis buffer (50 mM potassium phosphate at pH 8, 0).1 mM pyridoxal phosphate (Sigma P9255), 0.1x BugBuster (EMD Millipore Corp. 70921-4), 0.02 mg/mL bovine pancreas DNAse (GoldBio D-301-1), 1 mg/mL lysozyme (EMD Millipore Corp. 4403) for 1 hour at 37 °C. Lysates are clarified with centrifugation for 10 minutes (4,500 *g*, 4 °C) and stored at 4 °C until further use. All lysates are used on the same day as lysis. A reaction master mix composed of 625 μM indole (Sigma 13408) and 25 mM serine (Sigma S4500) is prepared from 0.5 M stocks in ethanol and KPi buffer (50 mM, pH 8), respectively. The master mix is warmed up to 37 °C prior to usage. The screens are set up by the addition of 160 μL of the pre-heated master mix to UV-transparent 96-well assay plates (Caplugs, catalog # 290-8120-0AF), and reactions are initiated by the addition of 40 μL lysate to each well. Plates are transferred to the plate reader (Tecan infinite M200 or Spark) and are shaken for 10 s. The absorbance is read at 290 nm for 30-45 min, or until tryptophan formation plateaus. The rate of tryptophan formation is determined by calculating absorbance change over time. In parallel, the sequences of all screened TrpB variants are determined using the 96-well plate starter cultures following the evSeq protocol.^59^ The inner primers sequences are detailed in **Table S4** of **Supporting Information**.

## Abbreviations

DeCOIL: Degenerate codon optimization for informed libraries
DC: Degenerate codon
DE: Directed evolution
ML: Machine learning
MLPE: Machine learning-assisted protein engineering
MLDE: Machine learning-assisted directed evolution
ftMLDE: Focused training machine-learning assisted directed evolution
GB1: Protein G subdomain B1
TrpB: β-subunit of tryptophan synthase from *Thermotoga maritima*
evSeq: Every variant sequencing

## Author Contributions

J.Y.: conceptualization, methodology, software, validation, investigation, writing–original draft, writing–review and editing, visualization, and funding acquisition. J.D.: data collection, investigation, writing–original draft, writing–review and editing, and funding acquisition. K.E.J.: methodology, data-collection, software, investigation, writing–original draft, writing–review and editing, and visualization. F-Z.L.: data-collection, software, writing–review and editing, and funding acquisition. Y.Y.: methodology, resources, writing–original draft, writing–review and editing, and funding acquisition. F.H.A.: resources, writing–original draft, writing–review and editing, and funding acquisition.

## Data Availability

All data needed to evaluate the conclusions in the paper are present in the paper and/or the Supporting Information.

## Acknowledgements

This material is based upon work supported by the U.S. Department of Energy, Office of Science, Office of Basic Energy Sciences, under Award Number DE-SC0022218. This report was prepared as an account of work sponsored by an agency of the United States Government. Neither the United States Government nor any agency thereof, nor any of their employees, makes any warranty, express or implied, or assumes any legal liability or responsibility for the accuracy, completeness, or usefulness of any information, apparatus, product, or process disclosed, or represents that its use would not infringe privately owned rights. Reference herein to any specific commercial product, process, or service by trade name, trademark, manufacturer, or otherwise does not necessarily constitute or imply its endorsement, recommendation, or favoring by the United States Government or any agency thereof. The views and opinions of authors expressed herein do not necessarily state or reflect those of the United States Government or any agency thereof. J.Y. and F.Z.L are supported by the National Science Foundation Graduate Research Fellowship. The authors thank Bruce Wittmann and Jennifer Sun for helpful discussions and Sabine Brinkmann-Chen for critical reading of the manuscript.

## Supporting Information

The supporting information is available free of charge.

## REFERENCES

(1) Romero, P. A.; Arnold, F. H. Exploring Protein Fitness Landscapes by Directed Evolution. 2009, 10 (DECEMbER). https://doi.org/10.1038/nrm2805.

(2) Smith, J. M. Natural Selection and the Concept of a Protein Space. Nature 1970, 225, 563–564.

(3) Arnold, F. H. Directed Evolution: Bringing New Chemistry to Life. Angew. Chem. Int. Ed. 2018, 57 (16), 4143–4148. https://doi.org/10.1002/anie.201708408.

(4) Yang, K. K.; Wu, Z.; Arnold, F. H. Machine-Learning-Guided Directed Evolution for Protein Engineering. Nat. Methods 2019, 16 (8), 687–694. https://doi.org/10.1038/s41592-019-0496-6.

(5) Wittmann, B. J.; Johnston, K. E.; Wu, Z.; Arnold, F. H. Advances in Machine Learning for Directed Evolution. Curr. Opin. Struct. Biol. 2021, 69, 11–18. https://doi.org/10.1016/j.sbi.2021.01.008.

(6) Freschlin, C. R.; Fahlberg, S. A.; Romero, P. A. Machine Learning to Navigate Fitness Landscapes for Protein Engineering. Curr. Opin. Biotechnol. 2022, 75, 102713. https://doi.org/10.1016/j.copbio.2022.102713.

(7) Hie, B. L.; Yang, K. K. Adaptive Machine Learning for Protein Engineering. Curr. Opin. Struct. Biol. 2022, 72, 145–152. https://doi.org/10.1016/j.sbi.2021.11.002.

(8) Ferguson, A. L.; Ranganathan, R. 100th Anniversary of Macromolecular Science Viewpoint: Data-Driven Protein Design. ACS Macro Lett. 2021, 10 (3), 327–340. https://doi.org/10.1021/acsmacrolett.0c00885.

(9) Mardikoraem, M.; Woldring, D. Machine Learning-Driven Protein Library Design: A Path Toward Smarter Libraries. In Yeast Surface Display; Traxlmayr, M. W., Ed.; Springer US: New York, NY, 2022; pp 87–104. https://doi.org/10.1007/978-1-0716-2285-8_5.

(10) Yu, T.; Boob, A. G.; Volk, M. J.; Liu, X.; Cui, H.; Zhao, H. Machine Learning-Enabled Retrobiosynthesis of Molecules. Nat. Catal. 2023. https://doi.org/10.1038/s41929-022-00909-w.

(11) Yu, T.; Cui, H.; Li, J. C.; Luo, Y.; Jiang, G.; Zhao, H. Enzyme Function Prediction Using Contrastive Learning. 2023.

(12) Sevgen, E.; Moller, J.; Lange, A.; Parker, J.; Quigley, S.; Mayer, J.; Srivastava, P.; Gayatri, S.; Hosfield, D.; Korshunova, M.; Livne, M.; Gill, M.; Ranganathan, R.; Costa, A. B.; Ferguson, A. L. ProT-VAE: Protein Transformer Variational AutoEncoder for Functional Protein Design; preprint; Synthetic Biology, 2023. https://doi.org/10.1101/2023.01.23.525232.

(13) Praljak, N.; Lian, X.; Ranganathan, R.; Ferguson, A. L. ProtWave-VAE: Integrating Autoregressive Sampling with Latent-Based Inference for Data-Driven Protein Design; preprint; Synthetic Biology, 2023. https://doi.org/10.1101/2023.04.23.537971.

(14) Bedbrook, C. N.; Yang, K. K.; Rice, A. J.; Gradinaru, V.; Arnold, F. H. Machine Learning to Design Integral Membrane Channelrhodopsins for Efficient Eukaryotic Expression and Plasma Membrane Localization. PLOS Comput. Biol. 2017, 13 (10), e1005786. https://doi.org/10.1371/journal.pcbi.1005786.

(15) Bedbrook, C. N.; Yang, K. K.; Robinson, J. E.; Mackey, E. D.; Gradinaru, V.; Arnold, F. H. Machine Learning-Guided Channelrhodopsin Engineering Enables Minimally Invasive Optogenetics. Nat. Methods 2019, 16 (11), 1176–1184. https://doi.org/10.1038/s41592-019-0583-8.

(16) Zhu, D.; Popova, G.; Nowakowski, T. J.; Schaffer, D. V. Optimal Trade-off Control in Machine Learning-Based Library Design, with Application to Adeno-Associated Virus (AAV) for Gene Therapy. 36.

(17) Bryant, D. H.; Bashir, A.; Sinai, S.; Jain, N. K.; Ogden, P. J.; Riley, P. F.; Church, G. M.; Colwell, L. J.; Kelsic, E. D. Deep Diversification of an AAV Capsid Protein by Machine Learning. Nat. Biotechnol. 2021, 39 (6), 691–696. https://doi.org/10.1038/s41587-020-00793-4.

(18) Mazurenko, S.; Prokop, Z.; Damborsky, J. Machine Learning in Enzyme Engineering. ACS Catal. 2020, 10 (2), 1210–1223. https://doi.org/10.1021/acscatal.9b04321.

(19) Greenhalgh, J. C.; Fahlberg, S. A.; Pfleger, B. F.; Romero, P. A. Machine Learning-Guided Acyl-ACP Reductase Engineering for Improved in Vivo Fatty Alcohol Production. Nat. Commun. 2021, 12 (1), 5825. https://doi.org/10.1038/s41467-021-25831-w.

(20) Xu, Y.; Verma, D.; Sheridan, R. P.; Liaw, A.; Ma, J.; Marshall, N. M.; McIntosh, J.; Sherer, E. C.; Svetnik, V.; Johnston, J. M. Deep Dive into Machine Learning Models for Protein Engineering. J. Chem. Inf. Model. 2020, 60 (6), 2773–2790. https://doi.org/10.1021/acs.jcim.0c00073.

(21) Miller, D. C.; Athavale, S. V.; Arnold, F. H. Combining Chemistry and Protein Engineering for New-to-Nature Biocatalysis. Nat. Synth. 2022, 1 (1), 18–23. https://doi.org/10.1038/s44160-021-00008-x.

(22) Qiu, Y.; Hu, J.; Wei, G.-W. Cluster Learning-Assisted Directed Evolution. Nat. Comput. Sci. 2021, 1 (12), 809–818. https://doi.org/10.1038/s43588-021-00168-y.

(23) Romero, P. A.; Krause, A.; Arnold, F. H. Navigating the Protein Fitness Landscape with Gaussian Processes. 2012. https://doi.org/10.1073/pnas.1215251110.

(24) Thomas, N.; Agarwala, A.; Belanger, D.; Song, Y. S.; Colwell, L. Tuned Fitness Landscapes for Benchmarking Model-Guided Protein Design; preprint; Synthetic Biology, 2022. https://doi.org/10.1101/2022.10.28.514293.

(25) Mardikoraem, M.; Woldring, D. Protein Fitness Prediction Is Impacted by the Interplay of Language Models, Ensemble Learning, and Sampling Methods; preprint; Bioinformatics, 2023. https://doi.org/10.1101/2023.02.09.527362.

(26) Aghazadeh, A.; Nisonoff, H.; Ocal, O.; Brookes, D. H.; Huang, Y.; Koyluoglu, O. O.; Listgarten, J.; Ramchandran, K. Epistatic Net Allows the Sparse Spectral Regularization of Deep Neural Networks for Inferring Fitness Functions. Nat. Commun. 2021, 12 (1), 5225. https://doi.org/10.1038/s41467-021-25371-3.

(27) Wu, Z.; Kan, S. B. J.; Lewis, R. D.; Wittmann, B. J.; Arnold, F. H. Machine Learning-Assisted Directed Protein Evolution with Combinatorial Libraries. Proc. Natl. Acad. Sci. 2019, 116 (18), 8852–8858. https://doi.org/10.1073/pnas.1901979116.

(28) Hopf, T. A.; Ingraham, J. B.; Poelwijk, F. J.; Schärfe, C. P. I.; Springer, M.; Sander, C.; Marks, D. S. Mutation Effects Predicted from Sequence Co-Variation. Nat. Biotechnol. 2017, 35 (2), 128–135. https://doi.org/10.1038/nbt.3769.

(29) Meier, J.; Rao, R.; Verkuil, R.; Liu, J.; Sercu, T.; Rives, A. Language Models Enable Zero-Shot Prediction of the Effects of Mutations on Protein Function; preprint; Synthetic Biology, 2021. https://doi.org/10.1101/2021.07.09.450648.

(30) Riesselman, A. J.; Ingraham, J. B.; Marks, D. S. Deep Generative Models of Genetic Variation Capture the Effects of Mutations. Nat. Methods 2018, 15 (10), 816–822. https://doi.org/10.1038/s41592-018-0138-4.

(31) Bloom, J. D.; Labthavikul, S. T.; Otey, C. R.; Arnold, F. H. Protein Stability Promotes Evolvability. Proc. Natl. Acad. Sci. 2006, 103 (15), 5869–5874. https://doi.org/10.1073/pnas.0510098103.

(32) Alley, E. C.; Khimulya, G.; Biswas, S.; AlQuraishi, M.; Church, G. M. Unified Rational Protein Engineering with Sequence-Based Deep Representation Learning. Nat. Methods 2019, 16 (12), 1315–1322. https://doi.org/10.1038/s41592-019-0598-1.

(33) Rao, R.; Bhattacharya, N.; Thomas, N.; Duan, Y.; Chen, X.; Canny, J.; Abbeel, P.; Song, Y. S. Evaluating Protein Transfer Learning with TAPE. 2019.

(34) Rao, R.; Liu, J.; Verkuil, R.; Meier, J.; Canny, J. F.; Abbeel, P.; Sercu, T.; Rives, A. MSA Transformer; 2021; p 2021.02.12.430858. https://doi.org/10.1101/2021.02.12.430858.

(35) Rives, A.; Meier, J.; Sercu, T.; Goyal, S.; Lin, Z.; Liu, J.; Guo, D.; Ott, M.; Zitnick, C. L.; Ma, J.; Fergus, R. Biological Structure and Function Emerge from Scaling Unsupervised Learning to 250 Million Protein Sequences. Proc. Natl. Acad. Sci. 2021, 118 (15). https://doi.org/10.1073/pnas.2016239118.

(36) Notin, P.; Dias, M.; Frazer, J.; Marchena-Hurtado, J.; Gomez, A.; Marks, D. S.; Gal, Y. Tranception: Protein Fitness Prediction with Autoregressive Transformers and Inference-Time Retrieval. arXiv May 27, 2022. http://arxiv.org/abs/2205.13760 (accessed 2022-05-30).

(37) Yang, K. K.; Zanichelli, N.; Yeh, H. Masked Inverse Folding with Sequence Transfer for Protein Representation Learning. 16.

(38) Yang, K. K.; Fusi, N.; Lu, A. X. Convolutions Are Competitive with Transformers for Protein Sequence Pretraining. 2022, 10.

(39) Hsu, C.; Nisonoff, H.; Fannjiang, C.; Listgarten, J. Learning Protein Fitness Models from Evolutionary and Assay-Labeled Data. Nat. Biotechnol. 2022, 1–9. https://doi.org/10.1038/s41587-021-01146-5.

(40) Hie, B. L.; Yang, K. K.; Kim, P. S. Evolutionary Velocity with Protein Language Models Predicts Evolutionary Dynamics of Diverse Proteins. Cell Syst. 2022. https://doi.org/10.1016/j.cels.2022.01.003.

(41) Nisonoff, H.; Wang, Y.; Listgarten, J. Augmenting Neural Networks with Priors on Function Values. 2022.

(42) Wittmann, B. J.; Yue, Y.; Arnold, F. H. Informed Training Set Design Enables Efficient Machine Learning-Assisted Directed Protein Evolution. Cell Syst. 2021, 1–20. https://doi.org/10.1016/j.cels.2021.07.008.

(43) Zhao, H.; Giver, L.; Shao, Z.; Affholter, J. A.; Arnold, F. H. Molecular Evolution by Staggered Extension Process (StEP) in Vitro Recombination. Nat. Biotechnol. 1998, 16 (3), 258–261. https://doi.org/10.1038/nbt0398-258.

(44) Packer, M. S.; Liu, D. R. Methods for the Directed Evolution of Proteins. Nat. Rev. Genet. 2015, 16 (7), 379–394. https://doi.org/10.1038/nrg3927.

(45) Voigt, C. A.; Martinez, C.; Wang, Z.-G.; Mayo, S. L.; Arnold, F. H. Protein Building Blocks Preserved by Recombination. Nat. Struct. Biol. 2002, 9 (7), 553–558. https://doi.org/10.1038/nsb805.

(46) Voigt, C. A.; Mayo, S. L.; Arnold, F. H.; Wang, Z.-G. Computational Method to Reduce the Search Space for Directed Protein Evolution. Proc. Natl. Acad. Sci. 2001, 98 (7), 3778–3783. https://doi.org/10.1073/pnas.051614498.

(47) Kille, S.; Acevedo-Rocha, C. G.; Parra, L. P.; Zhang, Z.-G.; Opperman, D. J.; Reetz, M. T.; Acevedo, J. P. Reducing Codon Redundancy and Screening Effort of Combinatorial Protein Libraries Created by Saturation Mutagenesis. ACS Synth. Biol. 2013, 2 (2), 83–92. https://doi.org/10.1021/sb300037w.

(48) Mena, M. A.; Daugherty, P. S. Automated Design of Degenerate Codon Libraries. Protein Eng. Des. Sel. 2005, 18 (12), 559–561. https://doi.org/10.1093/protein/gzi061.

(49) Jacobs, T. M.; Yumerefendi, H.; Kuhlman, B.; Leaver-Fay, A. SwiftLib: Rapid Degenerate-Codon-Library Optimization through Dynamic Programming. Nucleic Acids Res. 2015, 43 (5), e34–e34. https://doi.org/10.1093/nar/gku1323.

(50) Shimko, T. C.; Fordyce, P. M.; Orenstein, Y. DeCoDe: Degenerate Codon Design for Complete Protein-Coding DNA Libraries. Bioinformatics 2020, 36 (11), 3357–3364. https://doi.org/10.1093/bioinformatics/btaa162.

(51) Parker, A. S.; Griswold, K. E.; Bailey-Kellogg, C. Optimization of Combinatorial Mutagenesis. J. Comput. Biol. 2011, 18 (11), 1743–1756. https://doi.org/10.1089/cmb.2011.0152.

(52) Verma, D.; Grigoryan, G.; Bailey-Kellogg, C. Pareto Optimization of Combinatorial Mutagenesis Libraries. IEEE/ACM Trans. Comput. Biol. Bioinform. 2019, 16 (4), 1143–1153. https://doi.org/10.1109/TCBB.2018.2858794.

(53) Weinstein, E. N.; Amin, A. N.; Grathwohl, W.; Kassler, D.; Disset, J.; Marks, D. S. Optimal Design of Stochastic DNA Synthesis Protocols Based on Generative Sequence Models. bioRxiv October 29, 2021, p 2021.10.28.466307. https://doi.org/10.1101/2021.10.28.466307.

(54) Wu, N. C.; Dai, L.; Olson, C. A.; Lloyd-Smith, J. O.; Sun, R. Adaptation in Protein Fitness Landscapes Is Facilitated by Indirect Paths. eLife 2016, 5, e16965. https://doi.org/10.7554/eLife.16965.

(55) Protabit. https://triad.protabit.com/.

(56) Alford, R. F.; Leaver-Fay, A.; Jeliazkov, J. R.; O’Meara, M. J.; DiMaio, F. P.; Park, H.; Shapovalov, M. V.; Renfrew, P. D.; Mulligan, V. K.; Kappel, K.; Labonte, J. W.; Pacella, M. S.; Bonneau, R.; Bradley, P.; Dunbrack, R. L.; Das, R.; Baker, D.; Kuhlman, B.; Kortemme, T.; Gray, J. J. The Rosetta All-Atom Energy Function for Macromolecular Modeling and Design. J. Chem. Theory Comput. 2017, 13 (6), 3031–3048. https://doi.org/10.1021/acs.jctc.7b00125.

(57) Buller, A. R.; Brinkmann-Chen, S.; Romney, D. K.; Herger, M.; Murciano-Calles, J.; Arnold, F. H. Directed Evolution of the Tryptophan Synthase β-Subunit for Stand-Alone Function Recapitulates Allosteric Activation. Proc. Natl. Acad. Sci. 2015, 112 (47), 14599–14604. https://doi.org/10.1073/pnas.1516401112.

(58) Watkins-Dulaney, E.; Straathof, S.; Arnold, F. Tryptophan Synthase: Biocatalyst Extraordinaire. ChemBioChem 2021, 22 (1), 5–16. https://doi.org/10.1002/cbic.202000379.

(59) Wittmann, B. J.; Johnston, K. E.; Almhjell, P. J.; Arnold, F. H. evSeq: Cost-Effective Amplicon Sequencing of Every Variant in a Protein Library. ACS Synth. Biol. 2022, 11 (3), 1313–1324. https://doi.org/10.1021/acssynbio.1c00592.

(60) Wang, Z.; Combs, S. A.; Brand, R.; Calvo, M. R.; Xu, P.; Price, G.; Golovach, N.; Salawu, E. O.; Wise, C. J.; Ponnapalli, S. P.; Clark, P. M. LM-GVP: An Extensible Sequence and Structure Informed Deep Learning Framework for Protein Property Prediction. Sci. Rep. 2022, 12 (1), 6832. https://doi.org/10.1038/s41598-022-10775-y.

(61) El-Arini, K.; Veda, G.; Shahaf, D. Turning Down the Noise in the Blogosphere. 9.

(62) Khuller, S.; Moss, A.; Naor, J. (Seffi). The Budgeted Maximum Coverage Problem. Inf. Process. Lett. 1999, 70 (1), 39–45. https://doi.org/10.1016/S0020-0190(99)00031-9.

(63) Nemhauser, G. L.; Wolsey, L. A.; Fisher, M. L. An Analysis of Approximations for Maximizing Submodular Set Functions—I. Math. Program. 1978, 14 (1), 265–294. https://doi.org/10.1007/BF01588971.

(64) Chen, T.; Guestrin, C. XGBoost: A Scalable Tree Boosting System. In Proceedings of the 22nd ACM SIGKDD International Conference on Knowledge Discovery and Data Mining; 2016; pp 785–794. https://doi.org/10.1145/2939672.2939785.

(65) Gibson, D. G.; Young, L.; Chuang, R.-Y.; Venter, J. C.; Hutchison, C. A.; Smith, H. O. Enzymatic Assembly of DNA Molecules up to Several Hundred Kilobases. Nat. Methods 2009, 6 (5), 343–345. https://doi.org/10.1038/nmeth.1318.

