## Supporting Information for "DeCOIL: Optimization of Degenerate Codon Libraries for Machine Learning-Assisted Protein Engineering"

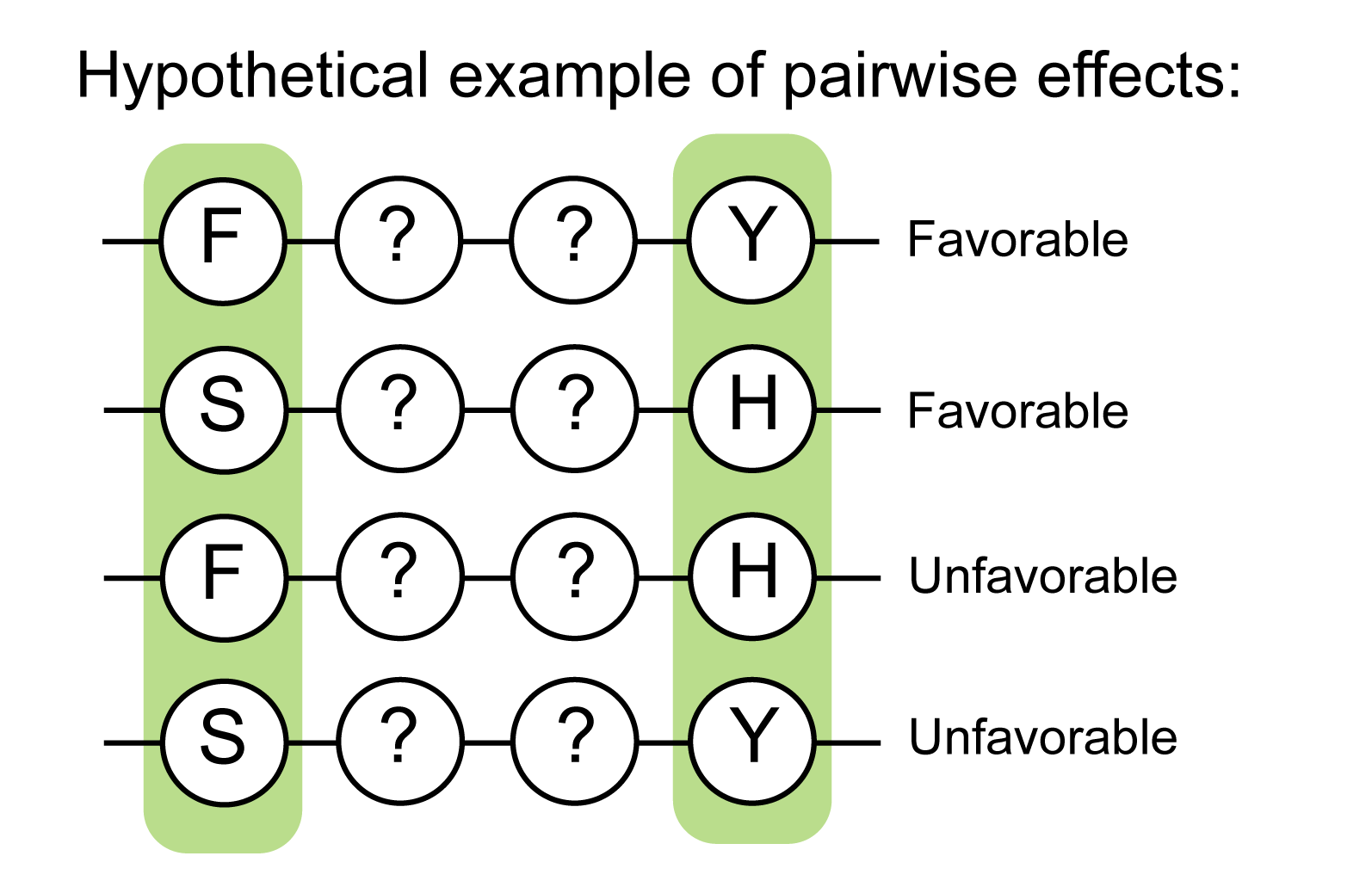

**Fig. S1. Hypothetical example of pairwise effects.** In this example, if the DC library frequently samples phenylalanine (F) at the first position, it should sample tyrosine (Y) at the fourth position, but not histidine (H), which is only favorable at the fourth position if serine (S) is at the first position.

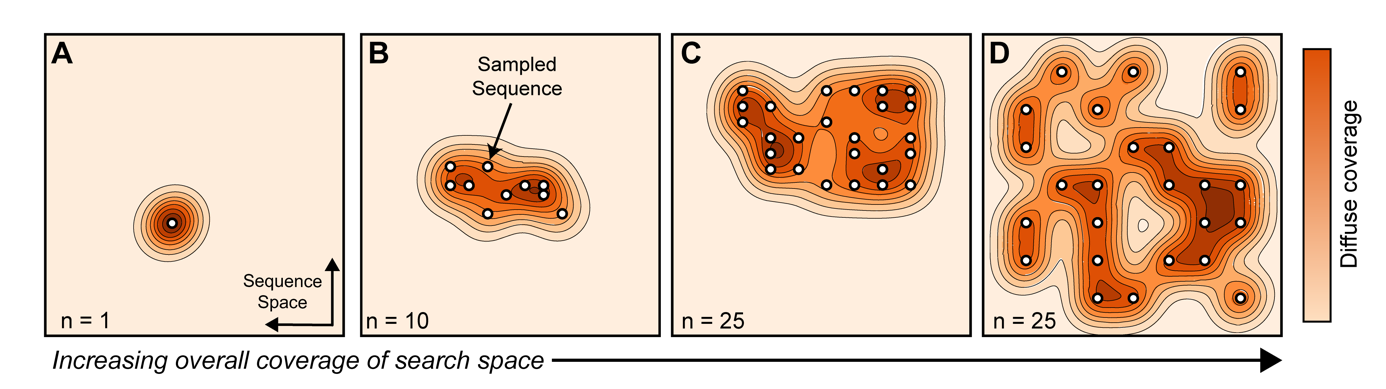

**Fig. S2. Examples of sets of amino acid sequences with varying degrees of overall unweighted coverage of the search space.**  Each sequence in the search space is assigned an individual coverage in one of two ways: (1) “step” coverage means that a sequence is assigned a coverage of 1 if it is sampled by a library and 0 otherwise, and (2) “diffuse” coverage means that a sequence is assigned an exponentially decaying coverage based on its distance to sequences already sampled by the library. Overall coverage is the sum of individual coverages for all sequences in the search space. Step coverage is directly proportional to the number of unique sequences, so the unweighted step coverage of the set of sequences portrayed in **(A)** is less than **(B)** and less than **(C)** and **(D),** but **(C)** and **(D**) have equal step coverage. By contrast, unweighted diffuse coverage increases from **(A)** to **(B)** to **(C)** to **(D)**. While step coverage and diffuse coverage are highly correlated, high step coverage is necessary but not sufficient for high diffuse coverage. Overall diffuse coverage is a more comprehensive metric for evaluating coverage of the search space because it accounts for amino acid types. Furthermore, it is generalizable to any sequence encoding.

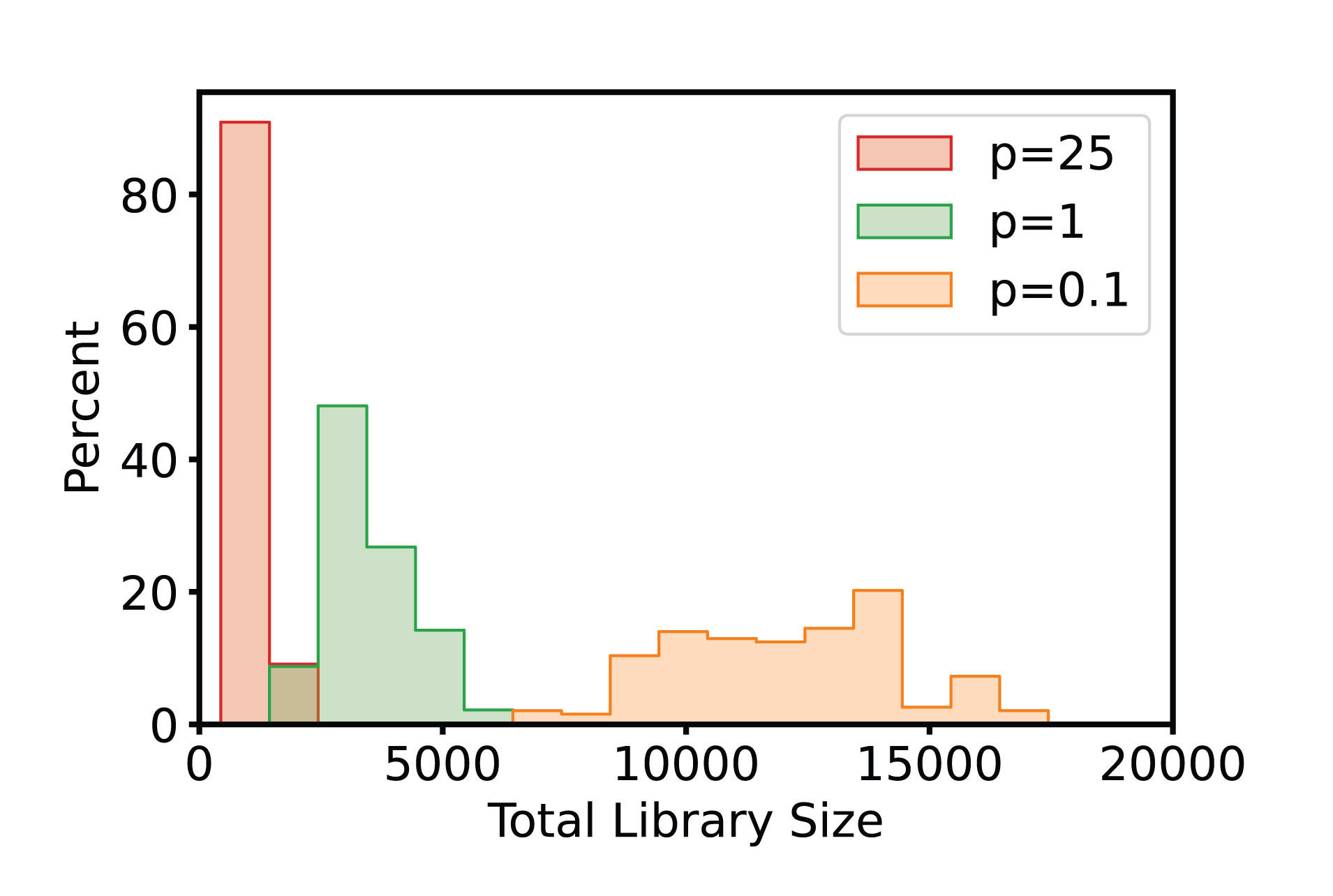

**Fig. S3. Histograms of total amino acid library sizes.** In general, DeCOIL-optimized libraries only cover a very small fraction of the total amino acid search space, which includes 160,000 variants for four sites. All optimized libraries use the Triad ∆∆G rank on the GB1 dataset with a screening size of 384. Figures in the Supporting Information can be reproduced from the codebase and results at https://github.com/jsunn-y/DeCOIL.

**Table S1. Performance of optimized DC libraries**.

|  | Mean Weighted Step Coverage  (*p* = 1) | Mean Fraction Unique | Mean Fraction Above 75^th^ Percentile of TRIAD ∆∆G | Mean Fraction Above 95^th^ Percentile of TRIAD ∆∆G | Median Mean True Fitness Sampled |
| --- | --- | --- | --- | --- | --- |
| NNK NNK NNK NNK | 0.44 | 0.88 | 0.25 | 0.06 | 0.009 |
| From DeCOIL: | | | | | |
| *p =* 0.1 (1 template) | 0.74±0.04 | 0.98±0.004 | 0.60±0.09 | 0.23±0.05 | 0.018±0.007 |
| *p =* 1 (1 template) | 0.85±0.03 | 0.93±0.02 | 0.86±0.05 | 0.46±0.11 | 0.036±0.014 |
| *p = 1* (2 templates) | 0.88±0.02 | 0.94±0.009 | 0.90±0.02 | 0.54±0.09 | 0.040±0.010 |
| *p = 1* (3 templates) | 0.89±0.01 | 0.95±0.007 | 0.92±0.02 | 0.56±0.08 | 0.041±0.009 |
| *p =* 25 (1 template) | 0.75±0.04 | 0.76±0.04 | 0.76±0.04 | 0.71±0.07 | 0.054±0.019 |
| From Swiftlib: | | | | | |
| Swiftlib1  NNS NNS KSC NNS (max size 1.6 x 10^6^) | 0.65 | 0.90 | 0.58 | 0.22 | 0.023 |
| Swiftlib2  NNS NNS GGA NNS (max size 4 x 10^5^) | 0.73 | 0.87 | 0.71 | 0.53 | 0.035 |
| Swiftlib3  DYA NNS GSA DNS (max size 1 x 10^5^) | 0.76 | 0.87 | 0.78 | 0.48 | 0.051 |

Optimized libraries using different values of *p* are skewed toward desired protein distributions, as measured by several metrics. Increasing the number of templates allowed during optimization increases the overall library performance slightly. On average, equivalent optimized DC libraries perform better than the top solution from Swiftlib. Swiftlib libraries are optimized to include the top 4000 sequences based on Triad ∆∆G, using different cutoffs for maximum library size. Mean fraction unique refers to the fraction of the screening library that is unique sequences, for sequences not containing stop codons. All DeCOIL-optimized libraries are based on the Triad ∆∆G rank on the GB1 dataset with a screening size of 384.

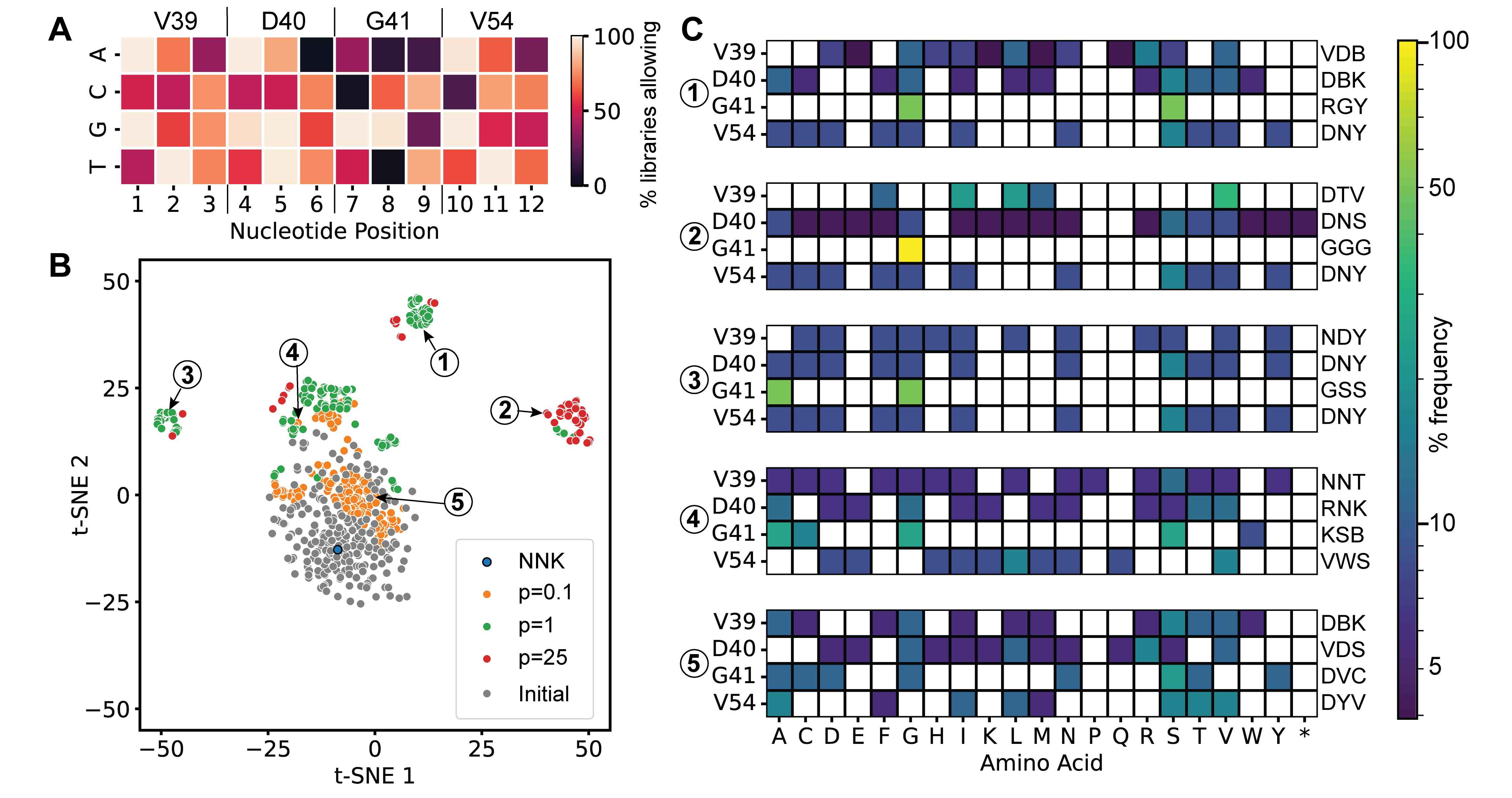

**Fig. S4. Optimized DC libraries are different from each other but show common learned patterns. (A)** Among the 240 DC libraries optimized using p=1, the percentages of them allowing each nucleotide (A, C, G, T) at each position. While most positions allow for highly diverse mixtures, certain positions strongly favor certain nucleotides, such as G over C at position 7 and G over T at position 8. **(B)** t-SNE visualization of optimized DC libraries, based on their corresponding amino acid distributions. Optimized libraries show several unique clusters, and libraries optimized using lower *p* values generally lie closer to the initial libraries (randomly generated as starting points for greedy hill climbing optimization). **(C)** Five selected libraries, defined by IUPAC nomenclature, and their corresponding amino acid distributions. While the amino acid distribution for residues 40 and 54 tend to vary less, there are noticeable differences in the amino acids sampled at residue 41. All optimized libraries use the Triad ∆∆G rank, on the GB1 dataset with a screening size of 384. An asterisk is used to denote stop codons.

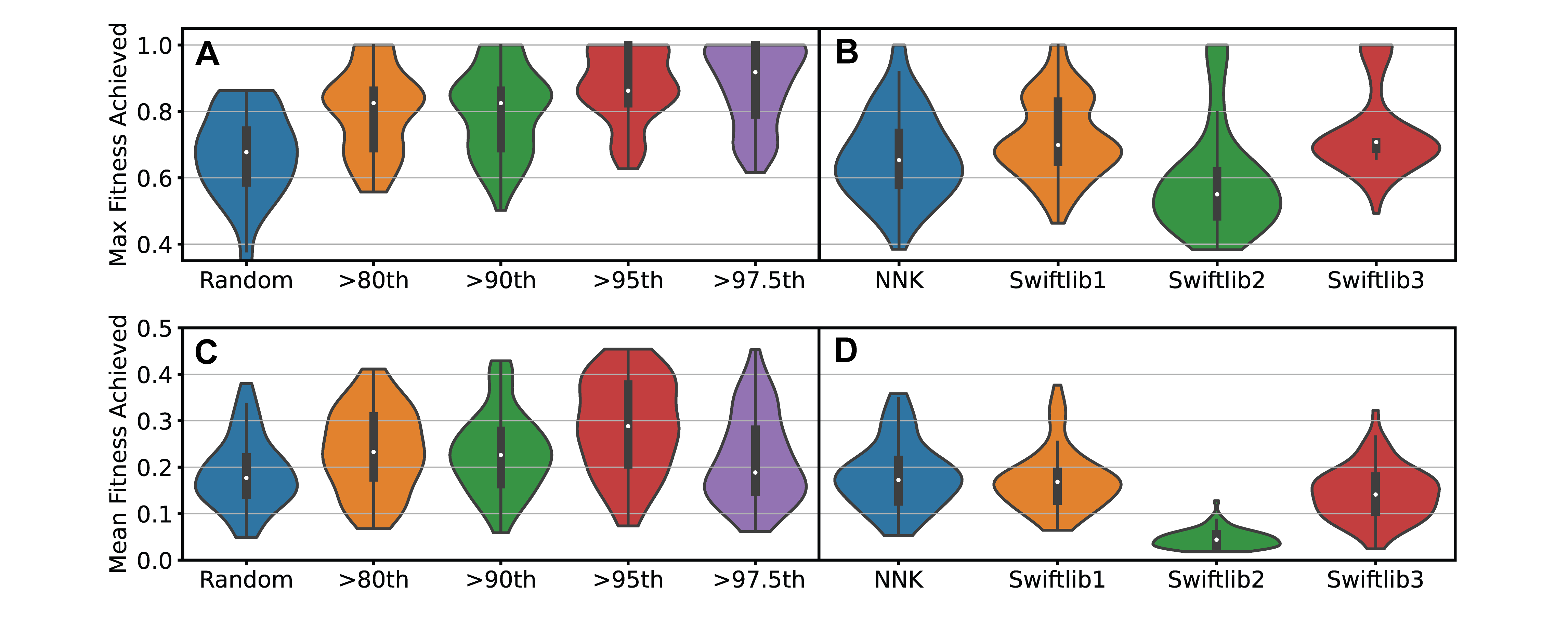

**Fig. S5. MLDE simulations with libraries that are treated as baselines. (A)** Max and **(C)** mean fitness achieved from downstream MLDE simulations using exact libraries as training sets. The exact libraries are sampled randomly from different focused training subsets of the search space. For example, >80^th^ means that variants are only sampled from the pool of variants with Triad ∆∆G above the 80^th^ percentile. **(B)** Max and (**D**) mean fitness achieved from downstream MLDE simulations using proteins sampled from several baseline DC libraries as training sets. Swiftlib libraries are defined in **Table S1**. Each ML model is an ensemble of five boosting models; training sets of 384 sequences are sampled from different libraries with 70 repeats; max and mean fitness are evaluated from the top 96 predictions of each ML model and normalized to the maximum fitness in the landscape. Tests are performed on the GB1 dataset, using the Triad ∆∆G rank as a ZS score.

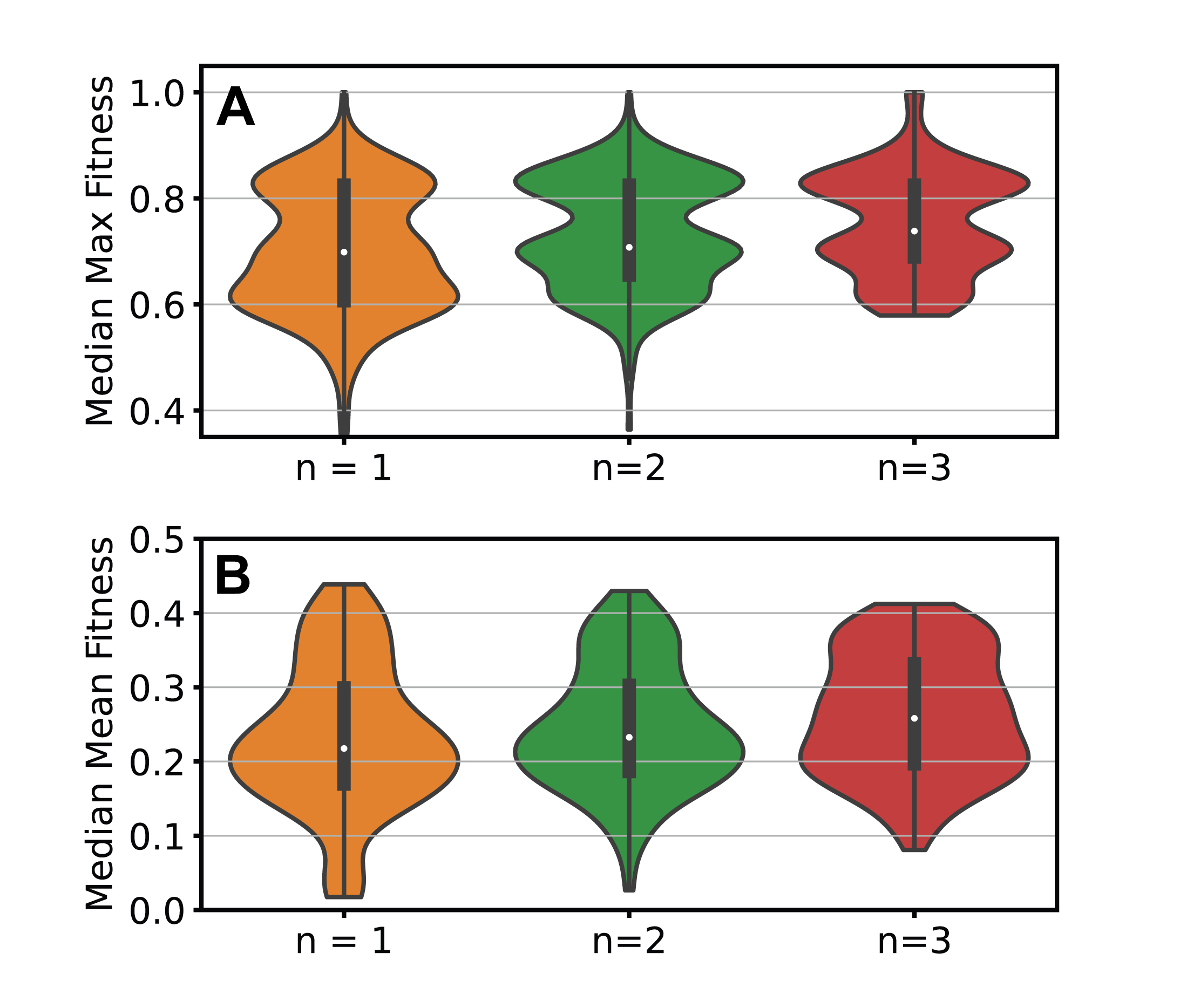

**Fig. S6. MLDE simulations of DC libraries optimized using increasing numbers of templates per library**. Median **(A)** max and **(B)** mean fitness achieved from downstream MLDE simulations, using proteins sampled from optimized DC libraries using p=1 as training sets. As the number of templates per library is increased, the performance of the average library increases slightly. Each ML model is an ensemble of 5 boosting models; training sets of 384 sequences (not all unique) are sampled from different libraries with 24 repeats; max and mean fitness are evaluated from the top 96 predictions of each ML model and normalized to the maximum fitness in the landscape. Tests are performed on the GB1 dataset, using the Triad ∆∆G rank as a ZS score.

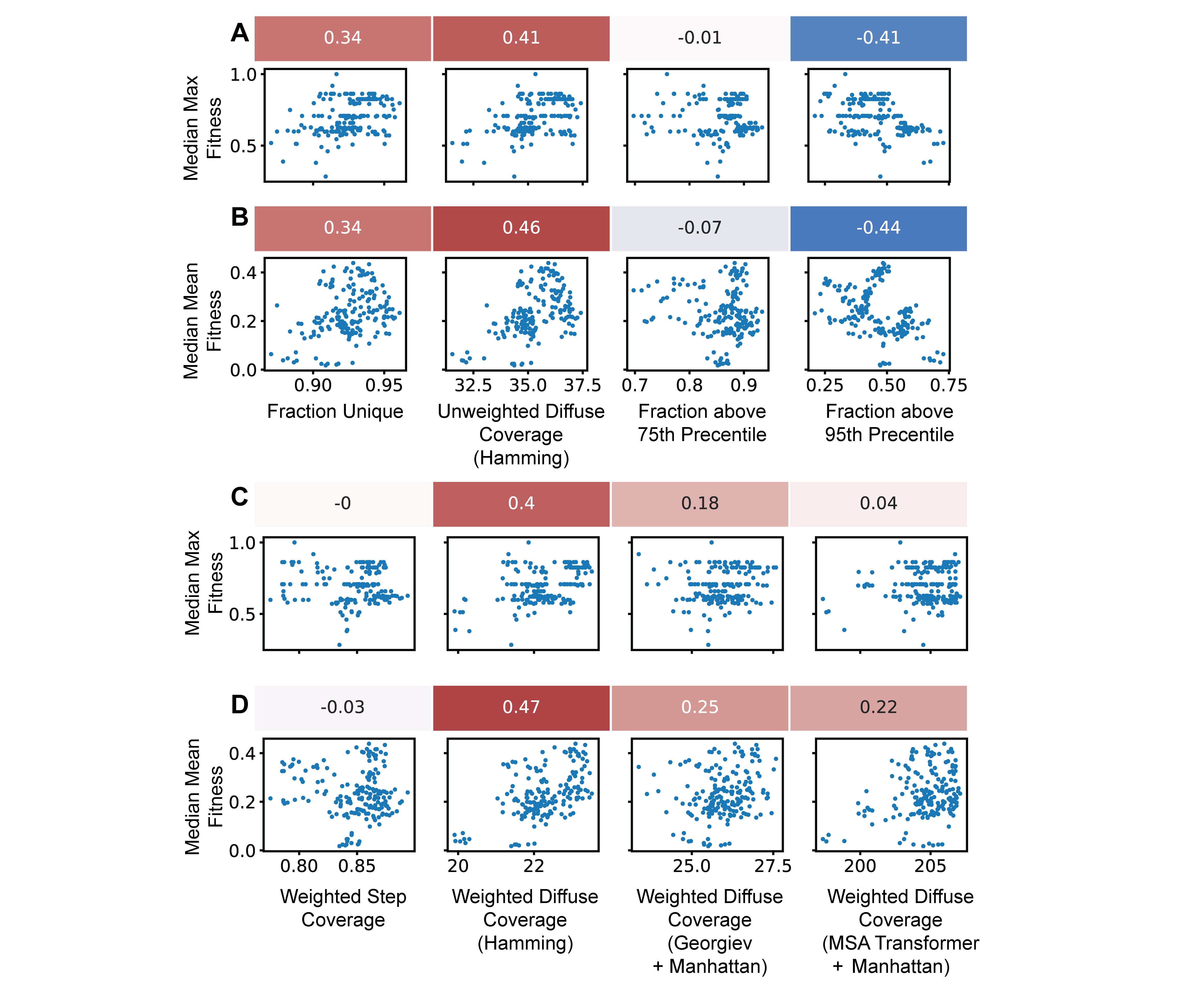

**Fig. S7. Correlations between downstream MLDE performance and various metrics.** Correlation between **(A)** max and **(B)** mean fitness achieved from downstream MLDE simulations and simplified metrics. The simplified metrics capture exploration or exploitation, but not necessarily both. Correlation between **(A)** max and **(B)** mean fitness achieved from downstream MLDE simulations and more robust metrics. The robust metrics capture exploration and exploitation. While unweighted and weighted diffuse coverage using Hamming distance are similarly strong predictors based on correlation, weighted diffuse coverage is better able to predict the libraries with the best MLDE performance. Each point represents a DeCOIL-optimized library using p=1. Correlations given are spearman ρ values. For diffuse coverage, σ=0.4, 20, and 0.01 are used for Hamming, Georgiev+Manhattan, and MSA transformer+Manhattan, respectively. All optimized libraries use the Triad ∆∆G rank, on the GB1 dataset with a screening size of 384.

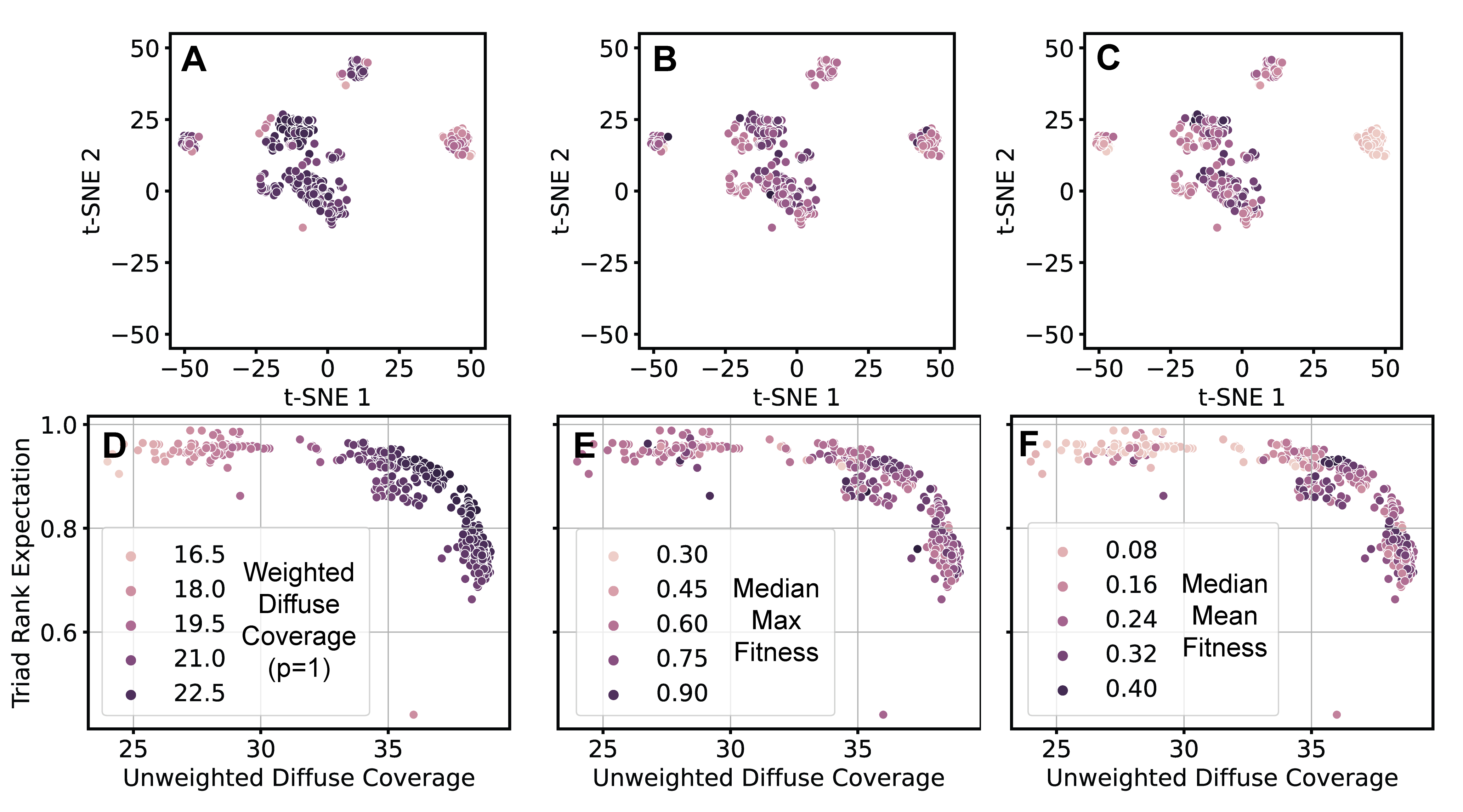

**Fig. S8. Visualization of DeCOIL-optimized libraries, colored by performance.** t-SNE visualization of optimized DC libraries, based on their corresponding amino acid distributions and colored by **(A)** weighted diffuse coverage, **(B)** median max fitness from downstream MLDE, and **(C)** median mean fitness from downstream MLDE. Exploration-exploitation tradeoff of optimized DC libraries colored by **(D)** weighted diffuse coverage, **(E)** median max fitness from downstream MLDE, and **(F)** median mean fitness from downstream MLDE. Diffuse coverage uses Hamming distance and σ = 0.4. All optimized libraries use the Triad ∆∆G rank, on the GB1 dataset with a screening size of 384.

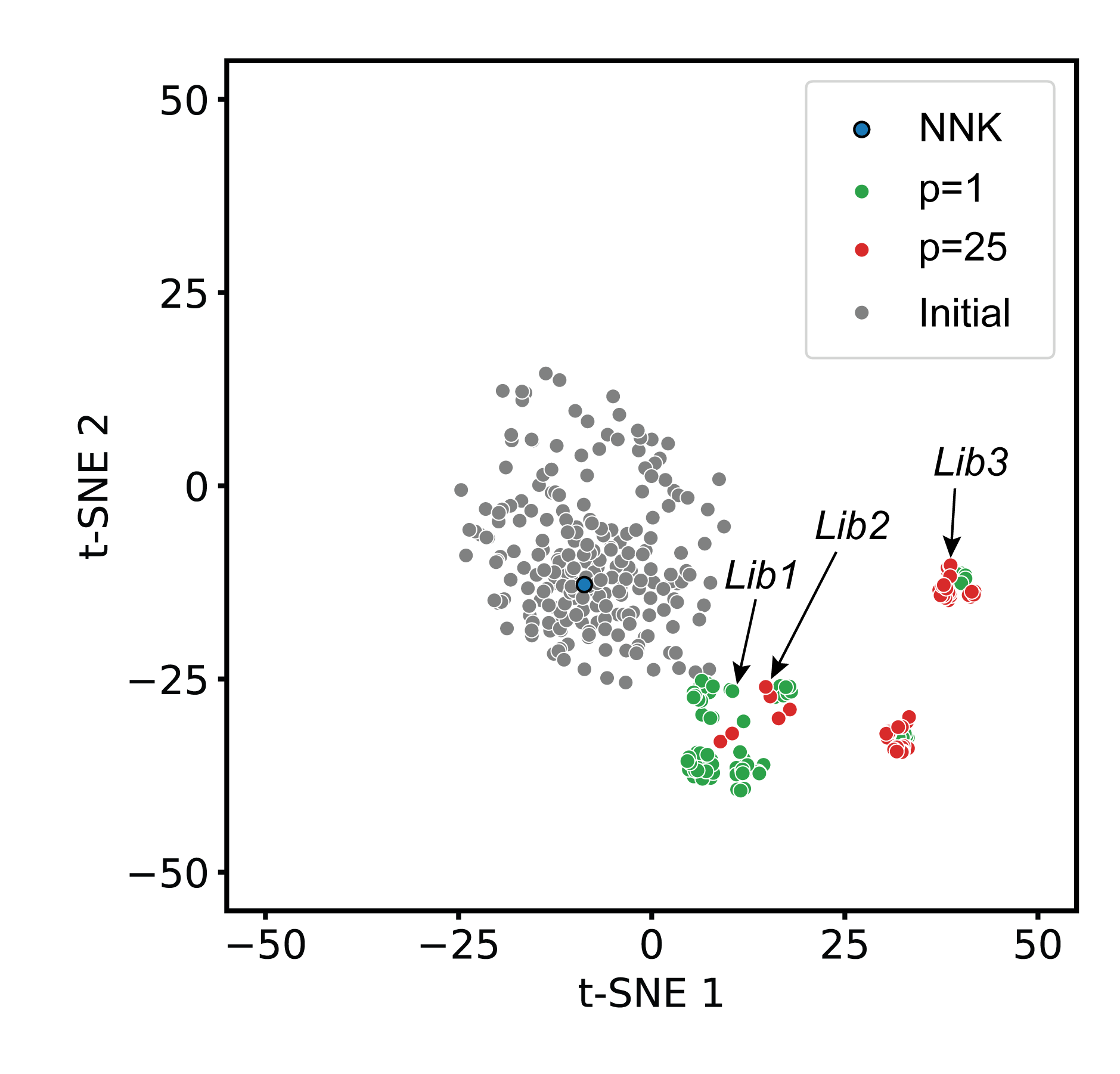

**Fig. S9.** **t-SNE visualization of optimized DC libraries, based on their corresponding amino acid distributions.** Optimization is performed on TrpB, using the EVmutation rank as a ZS score and a screening size of 384.

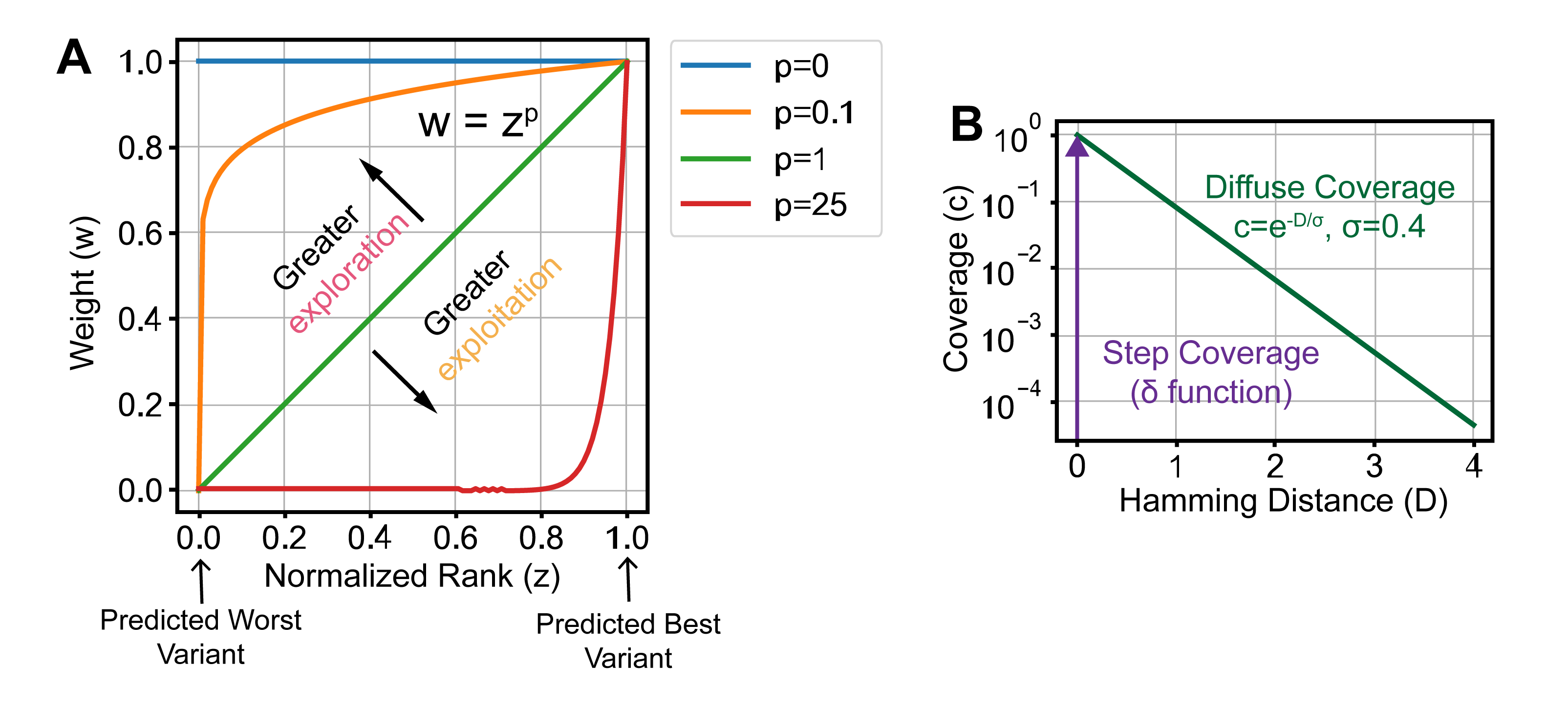

**Fig. S10.** **Graphs visualizing the equations used in the calculation of the optimization objective for DeCOIL.** DeCOIL optimizes an objective that aims to cover proteins variants with high weights. The *weight* factor is given as a power p of the ranking given from the predictor, thus increasing p will increase the amount of desired exploitation. If p=0, then the objective is unweighted (full exploration). A sequence in the search space is assigned a *coverage* factor in one of two ways: (1) the notion of “step” coverage means that a sequence is assigned a coverage of 1 if it sampled by a library and 0 otherwise and (2) the notion of “diffuse” coverage means that a sequence is assigned an exponentially decaying coverage based on its distance to sequences already sampled by the library.

**Table S2. Primer sequences for TrpB degenerate codon library construction.**

| **Name** | **Direction** | **Sequence** |
| --- | --- | --- |
| NNK_f | Forward | CCAACCTGCAGACCACCTAT**NNKNNKNNK**GGC**NNK**GTGGTTGGTCCGCATCCATATCC |
| Lib 1_f | Forward | CCAACCTGCAGACCACCTAT**YWYVNBDKS**GGC**DSY**GTGGTTGGTCCGCATCCATATCC |
| Lib 2_f | Forward | CCAACCTGCAGACCACCTAT**TWYNNBNYB**GGC**WCY**GTGGTTGGTCCGCATCCATATCC |
| Lib 3_f | Forward | CCAACCTGCAGACCACCTAT**YWYDBSNNB**GGC**TCT**GTGGTTGGTCCGCATCCATATCC |
| Gap_f | Forward | GTGGTTGGTCCGCATCC |
| AmpF | Forward | CCAACTTACTTCTGACAACGATCGGAGGACCGAAGGAGCTAACCGCTTTTTTGC |
| 188_r | Reverse | ATAGGTGGTCTGCAGGTTGGTAATCCAGTCACGCAGAGCTTCGT |
| AmpR | Reverse | CGATCGTTGTCAGAAGTAAGTTGGCCGCAGTGTTATCACTCATGGTTATGGCAG |

Beginning with a pET22b(+) backbone harboring a *Tm*9D8* gene (Figure S11), two fragments are amplified with a break in the ampicillin resistance gene and a gap where the variation will be introduced to reduce parent bias. Fragment A is constructed using the primers Gap_f and AmpR, and fragment B is generated using the primers AmpF and 188_r. Both fragments are amplified using the thermocycler methods showed in Table S3. The two fragments are incubated with 1 µL of DpnI (NEB R0176S) at 37 °C for 1 hour to digest the template plasmid. The products are then purified via gel electrophoresis and the Zymoclean Gel DNA Recovery Kit (Zymo Research D4002). Fragment A is next used as the template to generate fragment C, which contains the mutated region in the TrpB gene, using the primers NNK_f, lib1_f, lib2_f or lib3_f in combination with primer AmpR (**Table S3**, **Figure S12**). The degenerate codons inserted at position 182, 183, 184, and 186 of the TrpB gene are showed in bold. Fragment C is purified as above. All PCRs are conducted using the Phusion High-Fidelity PCR kit (NEB E0553). Fragments B and C are assembled using the Gibson assembly method. The four Gibson products are individually transformed to competent cells to generate the different TrpB degenerate codon libraries.

**Table S3. Thermal cycles used for the amplification of Fragment A, B and C for the construction of the TrpB degenerate codon libraries.**

| **Fragment A** | | |
| --- | --- | --- |
| Temperature (°C) | Time (sec) | Cycles |
| 98 | 00:30 | 1 |
| 98 | 00:10 | 5 |
| 55->59, 1C/s ramp | 00:15 |  |
| 72 | 1:00 |  |
| 98 | 00:10 | 25 |
| 59 | 00:15 |  |
| 72 | 1:00 |  |
| 72 | 10:00 | 1 |
| 10 | infinite | 1 |

| **Fragment B** | | |
| --- | --- | --- |
| Temperature (°C) | Time (sec) | Cycles |
| 98 | 00:30 | 1 |
| 98 | 00:10 | 30 |
| 72 | 00:15 |  |
| 72 | 2:45 |  |
| 72 | 10:00 | 1 |
| 10 | infinite | 1 |

| **Fragment C** | | |
| --- | --- | --- |
| Temperature (°C) | Time (sec) | Cycles |
| 98 | 00:30 | 1 |
| 98 | 00:10 | 5 |
| 58->63, 1C/s ramp | 00:15 |  |
| 72 | 1:00 |  |
| 98 | 00:10 | 25 |
| 63 | 00:15 |  |
| 72 | 1:00 |  |
| 72 | 10:00 | 1 |
| 10 | infinite | 1 |

ATGAAAGGCTACTTCGGTCCGTACGGTGGCCAGTACGTGCCGGAAATCCTGATGGGAGCTCTGGAAGAACTGGAAGCTGCGTACGAAGGAATCATGAAAGATGAGTCTTTCTGGAAAGAATTCAATGACCTGCTGCGCGATTATGCGGGTCGTCCGACTCCGCTGTACTTCGCACGTCGTCTGTCCGAAAAATACGGTGCTCGCGTATATCTGAAACGTGAAGACCTGCTGCATACTGGTGCGCATAAAATCAATAACGCTATCGGCCAGGTTCTGCTGGCAAAACTAATGGGCAAAACCCGTATCATTGCTGAAACGGGTGCTGGTCAGCACGGCGTAGCAACTGCTACCGCAGCAGCGCTGTTCGGTATGGAATGTGTAATCTATATGGGCGAAGAAGACACGATCCGCCAGAAACTAAACGTTGAACGTATGAAACTGCTGGGTGCTAAAGTTGTACCGGTAAAATCCGGTAGCCGTACCCTGAAAGACGCAATTGACGAAGCTCTGCGTGACTGGATTACCAACCTGCAGACCACCTATTACGTGTTCGGCTCTGTGGTTGGTCCGCATCCATATCCGATTATCGTACGTAACTTCCAAAAGGTTATCGGCGAAGAGACCAAAAAACAGATTCCAGAAAAAGAAGGCCGTCTGCCGGACTACATCGTTGCGTGCGTGAGCGGTGGTTCTAACGCTGCCGGTATCTTCTATCCGTTTATCGATTCTGGTGTGAAGCTGATCGGCGTAGAAGCCGGTGGCGAAGGTCTGGAAACCGGTAAACATGCGGCTTCTCTGCTGAAAGGTAAAATCGGCTACCTGCACGGTTCTAAGACGTTCGTTCTGCAGGATGACTGGGGTCAAGTTCAGGTGAGCCACTCCGTCTCCGCTGGCCTGGACTACTCCGGTGTCGGTCCGGAACACGCCTATTGGCGTGAGACCGGTAAAGTGCTGTACGATGCTGTGACCGATGAAGAAGCTCTGGACGCATTCATCGAACTGTCTCGCCTGGAAGGCATCATCCCAGCCCTGGAGTCTTCTCACGCACTGGCTTATCTGAAGAAGATCAACATCAAGGGTAAAGTTGTGGTGGTTAATCTGTCTGGTCGTGGTGACAAGGATCTGGAATCTGTACTGAACCACCCGTATGTTCGCGAACGCATCCGCCTCGAGCACCACCACCACCACCACTGA

**Figure S11. Tm9D8* parent sequence.**

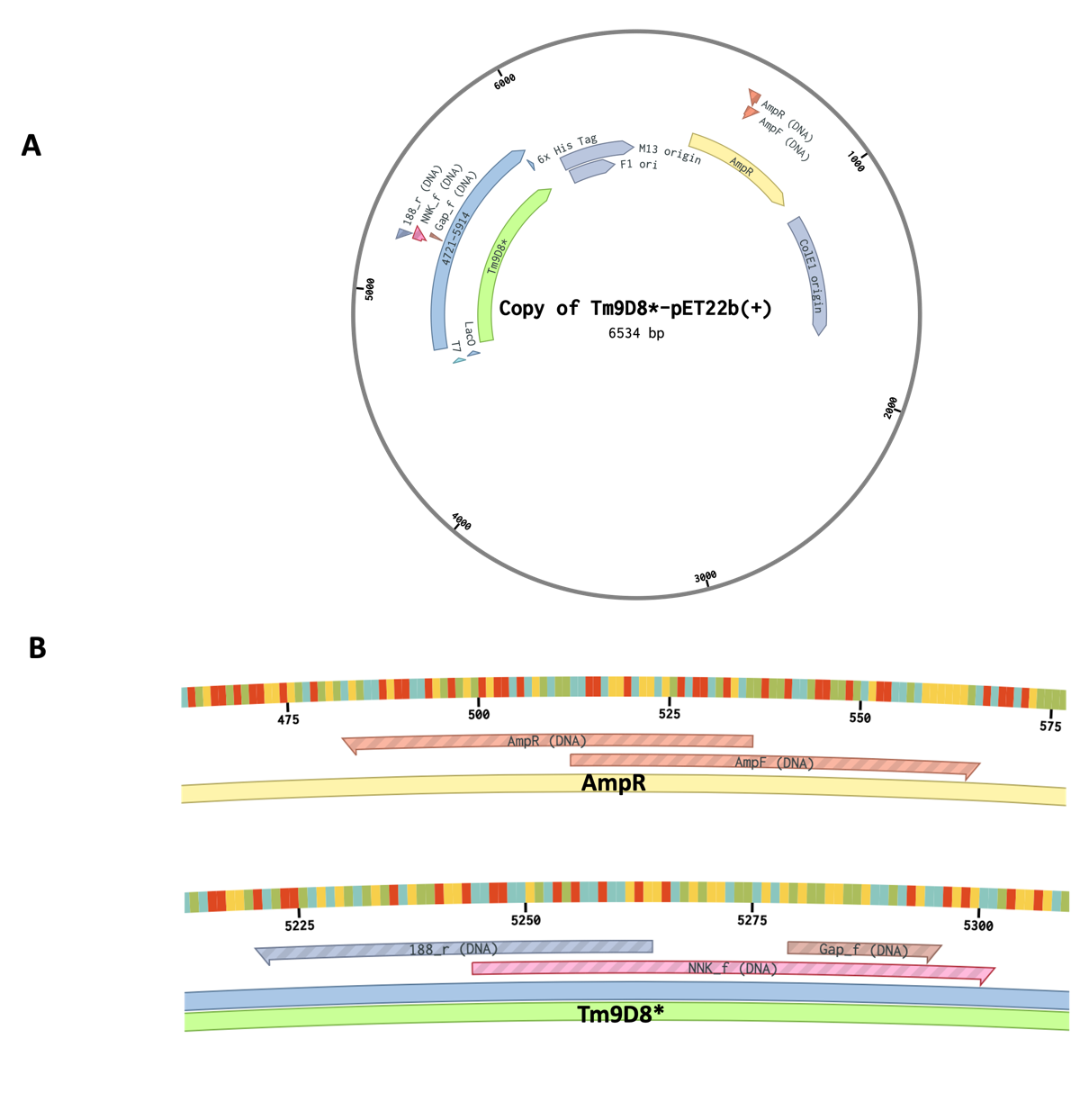

**Figure S12. Tm9D8*-pET22b(+) plasmid map. (A)** Circular map of the pET22b(+) plasmid displaying the position of the TrpB gene (Tm9D8*) and the ampicillin resistance cassette (AmpR). **(B)** Zoomed in section of the Tm9D8* and AmpR genes showing the position of the primers used for amplification of fragments A, B and C.

**Table S4. Inner primers used for evSeq of the TrpB degenerate codon libraries.**

| **Name** | **Direction** | **Sequence** |
| --- | --- | --- |
| 155_f | Forward | CACCCAAGACCACTCTCCGGCTGCTGGGTGCTAAAGTTGTACC |
| 215_r | Reverse | CGGTGTGCGAAGTAGGTGCTCCGGCAGACGGCCTTC |
